# Measuring instability in chronic human intracortical neural recordings towards stable, long-term brain-computer interfaces

**DOI:** 10.1101/2024.02.29.582733

**Authors:** Tsam Kiu Pun, Mona Khoshnevis, Thomas Hosman, Guy H. Wilson, Anastasia Kapitonava, Foram Kamdar, Jaimie M. Henderson, John D. Simeral, Carlos E. Vargas-Irwin, Matthew T. Harrison, Leigh R. Hochberg

**Affiliations:** Biomedical Engineering Graduate Program, School of Engineering, Brown University, Providence, RI, USA; School of Engineering, Brown University, Providence, RI, USA; Carney Institute for Brain Science, Brown University, Providence, RI, USA; Division of Applied Mathematics, Brown University, Providence, RI, USA; VA RR&D Center for Neurorestoration and Neurotechnology, Rehabilitation R&D Service, Providence VA Medical Center, Providence, RI, USA; Center for Neurotechnology and Neurorecovery, Dept. of Neurology, Massachusetts General Hospital, Boston, MA, USA; Department of Neuroscience, Brown University, Providence, RI, USA; Department of Neurology, Harvard Medical School, Boston, MA, USA; Department of Neurosurgery, Stanford University, Stanford, CA, USA; Wu Tsai Neurosciences Institute and Bio-X Institute, Stanford University, Stanford, CA, USA

**Author notes:** Co-senior authors (Matthew T. Harrison, Leigh R. Hochberg).

## Abstract

Intracortical brain-computer interfaces (iBCIs) enable people with tetraplegia to gain intuitive cursor control from movement intentions. To translate to practical use, iBCIs should provide reliable performance for extended periods of time. However, performance begins to degrade as the relationship between kinematic intention and recorded neural activity shifts compared to when the decoder was initially trained. In addition to developing decoders to better handle long-term instability, identifying when to recalibrate will also optimize performance. We propose a method to measure instability in neural data without needing to label user intentions. Longitudinal data were analyzed from two BrainGate2 participants with tetraplegia as they used fixed decoders to control a computer cursor spanning 142 days and 28 days, respectively. We demonstrate a measure of instability that correlates with changes in closed-loop cursor performance solely based on the recorded neural activity (Pearson *r* = 0.93 and 0.72, respectively). This result suggests a strategy to infer online iBCI performance from neural data alone and to determine when recalibration should take place for practical long-term use.

## Introduction

Intracortical brain-computer interfaces (iBCIs) have enabled people with tetraplegia to control external devices by decoding movement intentions from neural recordings ^1–7^. iBCIs can also restore communication by providing rapid point-and-click cursor control for applications such as typing, web browsing and navigating apps on a tablet ^8–13^, and can enable speech-to-text decoding for people with severe dysarthria ^14^. Decoders are typically trained during explicit calibration epochs that allow for simultaneous collection of recorded neural signals during instructed motor intentions ^6,15–17^. After training, decoding performance varies over time because of complex biological and device-related instabilities that are not fully understood ^18,19^. Persistent periods of decreased performance are commonly observed with existing decoding paradigms and remain one of the challenges that hinders wider adoption of iBCIs for people with paralysis ^12,18–22^. Restoration of good control after performance has degraded often requires the user to repeat a calibration task to retrain the decoder ^5,18^. Reducing the frequency and duration of explicit recalibration tasks is important for improving the utility of iBCIs. A step in this direction would be a method that could monitor performance and automatically determine when recalibration or other measures were necessary.

Here, we show that decoding performance on timescales of tens of seconds can be estimated from the same recorded neural signal used for motor decoding. The main idea is that persistent changes in performance likely result from statistical changes of some kind in the recorded neural signals. Measures of statistical changes might then be a good surrogate for measures of performance changes. We call this approach **MINDFUL** (**m**easuring **i**nstabilities in **n**eural **d**ata for **u**seful **l**ong-term iBCI). More specifically, given a target period for which average decoding performance is unknown, we calculate a statistical distance between the distribution of neural activity patterns during the target period and a similar distribution collected when performance was known to be good (such as when the decoder was first trained), as illustrated in **Fig. 1a**. The MINDFUL score obtained using Kullback-Leibler divergence (KLD) to compare neural activity patterns was found to correlate with decoding performance.

**Fig. 1.**
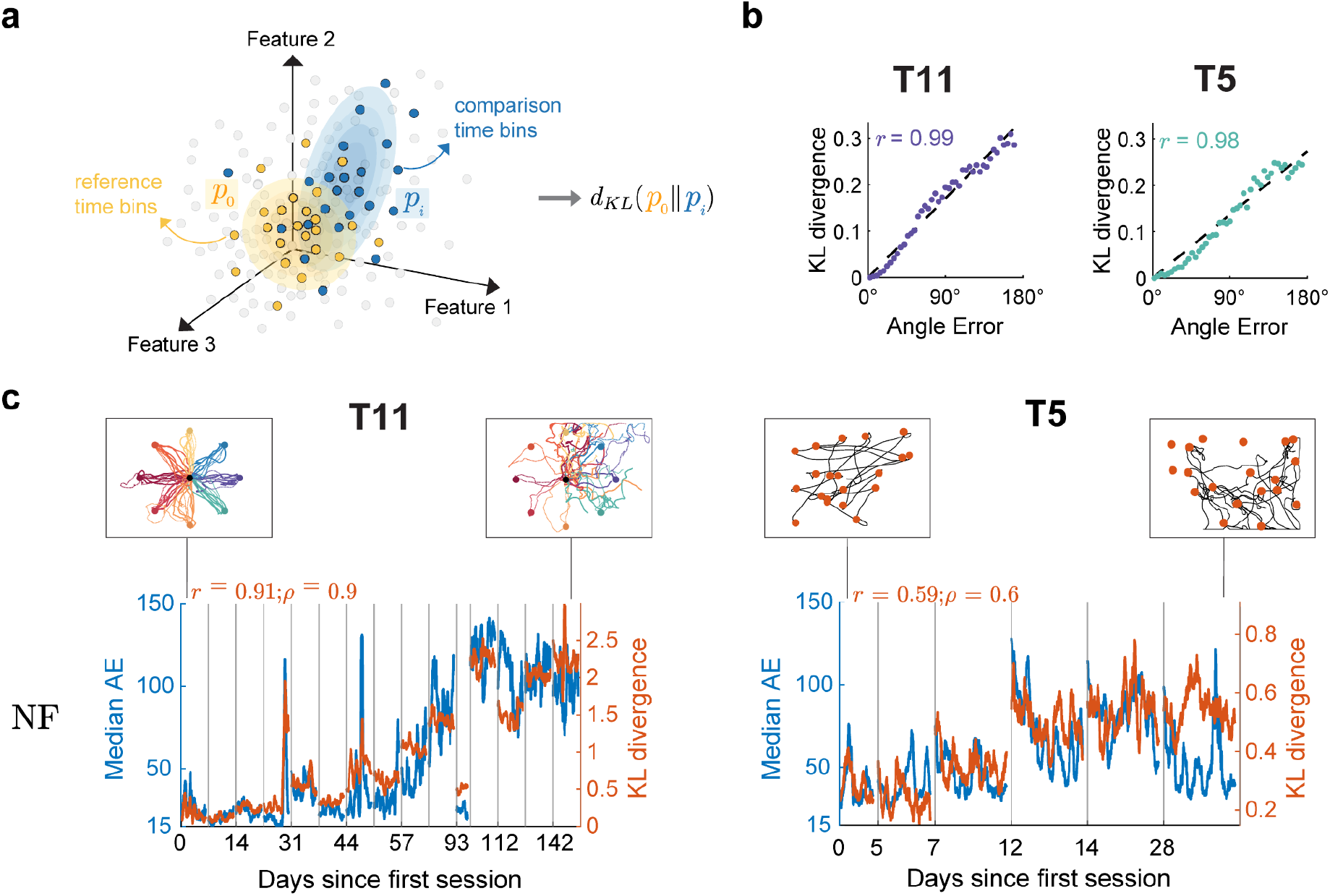
MINDFUL score correlates with performance over time. **(a)** Illustration of how the MINDFUL metric is calculated. Each dot symbolizes neural features at a given time bin in any given session, colored by whether the time bin is included for estimating the reference distribution (yellow), or for comparison (blue), or neither (gray). The difference in distributions is quantified by Kullback-Leibler divergence (KLD) between the reference distribution, *p*0and the comparison distribution,*pi*. **(b)** Binned samples of neural features were grouped according to decoder performance (AE), including data from all sessions. A total of 45 different distributions were generated, with AE increasing in 4° intervals from 0° to 180°. Bins with low AE (< 4°) were chosen as the reference distribution, and compared against the other 44 for T11 (left panel) and T5 (right panel). The dotted line which represents the best linear regression fit, along with the Pearson correlation coefficients,*r*, is shown. **(c)**The reference distribution was estimated from neural features (NF) time bins where AE < 4°, limited to day 0 where the decoder was first deployed for T11, and day 0 and 5 for T5. For subsequent sessions, neural distributions for comparison were constructed using an overlapping sliding window of 60 seconds at 1 second intervals. The KLD (right y-axis) is overlaid onto median AE (left y-axis in blue) across all recorded sessions for T11 (left panel) and T5 (right panel). Gray lines indicate the beginning of the session. Pearson and Spearman rank correlation, *r* and *p* respectively, quantify the relationship between the KLD and median AE. Insets present examples of cursor control of the task in the first and last session. For T11, cursor trajectories for all trials during a 5-min block are shown. Each color represents a peripheral target in a center-out-and-back task. For T5, cursor trajectories of the first 20 trials of a block are shown, along with the corresponding target presented at a random location on the screen in each trial.

Changes in decoding performance can be attributed to many types of variability in the recorded neural signals. BCI decoding algorithms typically model rapid fluctuations in neural features (on timescales of tens of milliseconds) that have no apparent correlation with motor intention as stochastic *noise*. Noise is a useful explanation for why decoding performance changes from instant to instant, but it generally does not account for persistent changes in (average) decoding performance that last seconds or more. We ascribe such persistent changes to *model drift*, i.e., changes in the relationship between recorded neural signals and motor intention that are not accounted for by the decoding paradigm. Model shift, dataset shift, and nonstationarity are terms that have been used synonymously in the literature for this type of phenomenon ^23–30^. Model drift can be attributed to various factors such as changes in action potential waveforms ^18,31,32^, neural tuning profiles ^33–35^, cognitive strategy or plasticity due to learning ^36–38^, material degradation and tissue responses to the recording device ^39,40^, and array micro-movements ^41^. The type and magnitude of model drift results in various forms of performance degradation ^19^, sometimes necessitating decoder recalibration to restore control. Existing solutions to reduce the need for recalibration tasks include adaptive decoders that require shorter recalibration sessions to maintain or restore stable performance ^3,10,42^, self-supervised recalibration using retrospective labeling that avoids explicit recalibration sessions ^26,27,43,44^, and robust decoders that experience less model drift by extracting stable, time-invariant features from high-dimensional recordings ^20–22,45–51^ or by adaptively adjusting decoder parameters ^12,52^.

The model drift that influences performance is necessarily a property of the joint distribution of recorded neural signals and motor intention. It is not *a priori* clear, however, that model drift related to performance can be meaningfully identified from the recorded neural signals alone, which is what MINDFUL attempts to capture. MINDFUL differs from previously studied statistical tests for model drift that additionally require knowledge of movement intention ^28,53,54^. Since true movement intention is often unavailable in iBCI applications where people with paralysis control an external device without being cued to acquire targets (e.g., a cursor on a tablet computer being used to send an email^22^), an approach like MINDFUL based only on the (marginal) distribution of recorded neural activity is much more widely applicable.

Here, we present and validate an approach to predict closed-loop decoding performance without the knowledge of true movement intention. MINDFUL was applied on longitudinal datasets where performance changed over long time periods as two people with tetraplegia, designated as T11 and T5, were using an iBCI. Each participant used an iBCI to control a computer cursor to perform target acquisition tasks on a screen. Target acquisition tasks permit observation of (presumed) motor intention and, hence, can be used to directly measure decoding performance. The kinematic decoders were held fixed across all sessions so that persistent changes in performance could be ascribed to model drift and not to changes associated with the decoder. Briefly, MINDFUL represents changes in neural distributions relative to a reference distribution where the decoder was initially applied. MINDFUL is solely based on the recorded neural activity, without requiring information about the target locations. The resulting MINDFUL score was highly correlated with changes in closed-loop cursor performance over time.

## Results

### Fixed decoders result in initially stable and then unstable performance across month-long sessions

To first establish a baseline for decoder performance, we deployed fixed decoders ^27,51^ for the purpose of identifying, over a comparatively long period, how neural instabilities may lead to deteriorating control. Data were collected from 15 consecutive research sessions spanning 142 days of T11 performing a center-out-and-back task using a fixed nonlinear (recurrent neural network) decoder, as previously described ^51^ (see **Methods**). As part of another study ^27^, T5 performed a random target task for six consecutive sessions spanning 28 days using a fixed linear decoder (see **Methods**). To quantify closed-loop cursor performance, *angle error* (AE) between the inferred intended directional vector (cursor-to-target position) and the decoded velocity vector was used (see **Methods**). AE is a valuable metric for capturing performance as it is sensitive to instantaneous cursor direction change, and can be averaged across any range of time. T11 achieved stable, high performance online cursor control for the first three months. The median AE per trial for T11 for sessions during the first three months was lower than later sessions on average (trial day 658-751: 26.8°±22.6°; trial day 758-800: 88.4°±46.1°; p < 0.001; Wilcoxon rank sum). For T5, the first three sessions demonstrated lower AE than the later three sessions (trial day 2121-2128: 39.6°±23.9°; trial day 2133-2149: 58.8°±31.7°; p < 0.001, Wilcoxon rank sum). Brief recovery from decrease in performance was observed in both participants (93 days after the initial session for T11 and 28 days after the initial session for T5), indicating fixed decoders may not necessarily result in a steady decline in cursor control over time (**Supplementary Fig. 1)**.

### Comparing distributions of neural activity patterns

MINDFUL is based on comparing the distribution of recorded neural activity patterns in a target dataset (usually with unknown decoding performance) to a similar distribution in a reference dataset (usually with known and good decoding performance). **Fig. 1a** provides an illustration. The choices of neural features and measure of statistical dissimilarity are important for practical use. Here we used a measure of statistical dissimilarity based on the well-known Kullback-Leibler divergence (KLD). In principle other measures of dissimilarity could be used (see **Methods**). Neural features were all derived from the underlying inputs to the kinematic decoder: threshold-crossing spike rate and spike power for T11 and spike rate only for T5, extracted in 20 ms non-overlapping bins (see **Methods**).

### Statistical distance between neural activity patterns correlates with performance

Having established datasets where fixed decoders result in periods of both stable and fluctuating closed-loop performance, we first investigated the underlying premise of MINDFUL that the distribution of neural activity patterns varies systematically with decoder performance. First, neural features pooled from all sessions were categorized into groups based on performance in terms of instantaneous (20 ms) AE. KLD was computed to assess the differences in neural feature distribution at instances with low AE (< 4°) to other distributions at instances with varying levels of AE (see **Methods**). For both participants, the relationship between the KLD and performance was found to be remarkably linear and strongly correlated (T11: Pearson *r* = 0.985, *p* < 10^−33^; T5: Pearson *r* = 0.983, *p* < 10^−31^; see **Fig. 1b**). Neural feature distributions of instances with low AE demonstrated high similarity (lower KLD) to the reference distribution of neural features from low AE instances, and the KLD increased linearly as the compared neural feature distributions were drawn from instances with larger AE.

This is a proof-of-concept that statistical distance between distributions of neural activity patterns can correlate strongly with decoding performance. It does not, however, provide a measure of performance that would be useful in a clinical setting for detecting persistent changes in decoding performance that might arise from model drift. Instantaneous AE must be known a priori to define the collections of neural features that are compared in each point in **Fig. 1b** and MINDFUL is designed to be used in situations where AE is not known, at least not for the target distribution. Moreover, AE can be a result of noise (transient variability) or model drift (persistent changes) or both. **Fig. 1b** does not distinguish among these even though model drift is the phenomenon of interest here. The linear relationship observed in **Fig. 1b** can be recreated in simulation using noise or using model drift or using both (see **Supplementary Fig. 2**).

### MINDFUL correlates with decoding performance

Towards developing a predictor of decoder performance based on neural activity for online application, we define a measure called the MINDFUL score to study the effect of drifts that persist over timescales relevant to chronic continuous iBCI use. Using the same concept as illustrated in **Fig. 1a**, but instead of grouping by AE as in **Fig. 1b**, neural feature distributions were estimated from collections of time bins aggregated using a 60-second sliding window, regardless of the performance during that window. The reference distribution is also estimated from instances of low AE as in Fig. 1b, but it is sub-selected from only the initial session(s) when the decoder is first deployed. As time progresses, we update the MINDFUL score which is based on the KLD between the reference distribution and the subsequent neural feature distributions from the sliding window (see **Methods**). To validate this method, MINDFUL score is correlated against the median AE calculated in the same 60-second sliding intervals as in estimating the neural distributions. Strong correlations were found between MINDFUL score and median AE across sessions, especially for T11 (See **Fig. 1c**. T11: Pearson *r* = 0.91, *p* < 0.001; Spearman ρ = 0.90, *p* < 0.001; T5: Pearson *r* = 0.59, *p* < 0.001; Spearman ρ = 0.60, *p* < 0.001; see **Methods**). This suggests that MINDFUL can be a viable measure for tracking performance in real-time, as the statistical properties of neural features aggregated over a longer timescale window reflects information about how the decoder performance drifts over time without needing to know performance (AE) in advance.

### Correlation to performance increases by combining neural features and decoder outputs

The neural features (NF) in **Fig. 1b-c** were derived from principal components analysis (PCA; see **Methods**) and need not reflect the sources of variability most predictive of decoder performance. Using features more closely related to the decoder output might strengthen the relationship between the MINDFUL score and AE. One such feature is the output from the decoder itself – in this case, the predicted 2-dimensional velocity vector,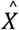. Note that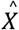 alone cannot be used to define AE. It is the relationship Between 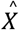 and the true intended direction that defines AE. Nevertheless, changes in the distribution of 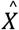 over a time interval may reflect changes in decoder performance. Likewise to **Fig. 1c**, a 60-second sliding window when estimating the mean and the variance of the distributions. Using only the 2-dimensional feature 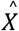 showed reduced correlations between the MINDFUL score and AE than from just NF for T11, but increased for T5 (**Fig. 2a**. T11: Pearson *r* = 0.464, *p* < 0.001; Spearman ρ = 0.407, *p* < 0.001; T5: Pearson *r* = 0.704, *p* < 0.001; Spearman ρ = 0.763, *p* < 0.001). When using the 4-dimensional feature created by concatenating 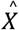 and 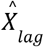, where 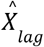 comes from the previous time bin of 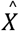 (20ms earlier), there was an increased correlation to AE for both T11 and T5 (**Fig. 2b**. T11: Pearson *r* = 0.819, *p* < 0.001; Spearman ρ = 0.840, *p* < 0.001; T5: Pearson *r* = 0.702, *p* < 0.001; Spearman ρ = 0.765, *p* < 0.001). Lastly, the combination of inputs and outputs of the decoder, i.e. the low-dimensional NF, 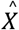, and 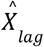 resulted in the highest correlation between KLD and AE for both participants (**Fig. 2c**. T11: Pearson *r* = 0.926, *p* < 0.001, Spearman ρ = 0.913, *p* < 0.001, T5: Pearson *r* = 0.719, *p* < 0.001, Spearman ρ = 0.759, *p* < 0.001).

**Fig. 2.**
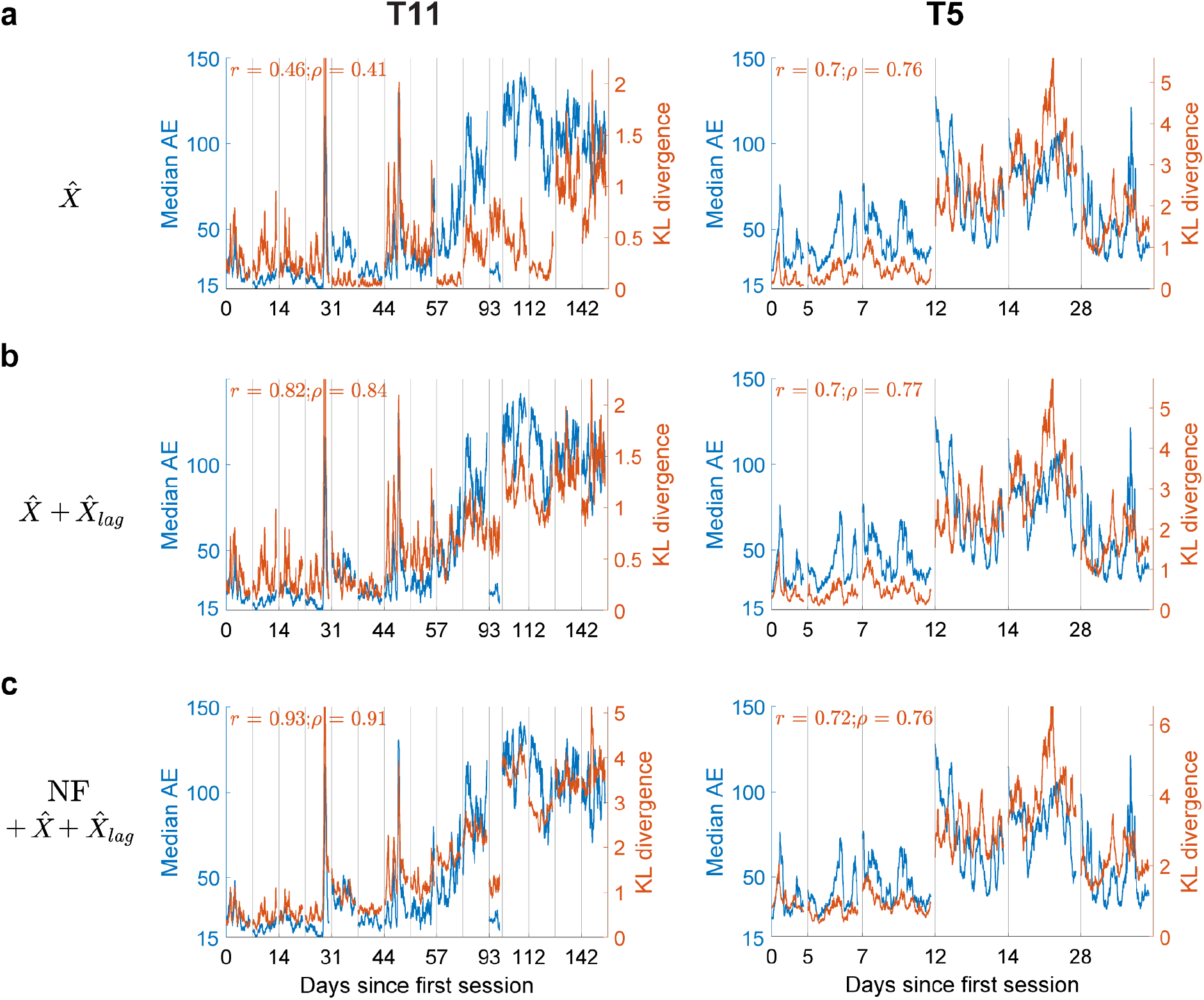
Incorporating decoder outputs in the MINDFUL score maintains high correlation with performance. **(a)** The KLD (right y-axis) between distributions of decoded directional vectors, 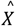, with respect to the first session overlaid onto median AE (left y-axis in blue) across all recorded sessions for T11 (left panel) and T5 (right panel). Subsequent neural distributions and median AE were updated every 1 second over a 60-second sliding window. Pearson *r*, and Spearman rank correlation coefficients ρ, between KLD and median AE are shown. **(b)** The KLD between distributions of 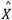 and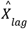overlaid onto median AE. **(c)** The KLD between distributions of the combination of derived neural features (as shown in **Fig. 1c**), decoded directional vectors, 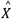, and 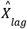, overlaid onto median AE.

### The MINDFUL score reflects changes in feature tuning

We next investigated the types of changes in neural data captured by the MINDFUL score. Changes in directional tuning have been shown to reduce performance in both online and offline BCI studies ^10,12,18–22^. Directional tuning was quantified by fitting a cosine function to normalized neural features to obtain estimates of preferred direction (PD) and modulation depth (MD) ^33,55–57^. 154 out of 384 features and 85 out of 192 features, for T11 and T5 respectively, had significant directional tuning for at least half of all recording sessions (F-test, *p* < 0.05, see **Methods**). Changes in tuning in these features were tracked over time (see **Methods**). (Although our analysis here is focused on the tuning of *individua*l features, in the supplement we use a different approach to show that changes in the distribution of *population* activity are statistically significant; see **supplementary Fig 3**.)

Tuning properties shifted gradually for the majority of features in T11 (**Fig. 3a-b**). 125 out of 151 tuned features exhibited significant change in both MD and PD ^35^ in at least one session (see **Methods**). Fitted tuning curves across sessions for two example features illustrated changes in modulation depth and modulation lost, respectively for T11 (**Fig 3c**). The average change in PD across these features was larger on days where performance was worse (day 7-93: 46.8°±31.2°; day 100-142: 62.4°±34.3°; *p* < 10^−7^ Wilcoxon rank sum). Average absolute change in MD in later sessions was also found to be significantly larger (day 7-93: 0.107±0.067; day 100-142: 0.159±0.129; *p* < 10^−8^, Wilcoxon rank sum).

**Fig. 3.**
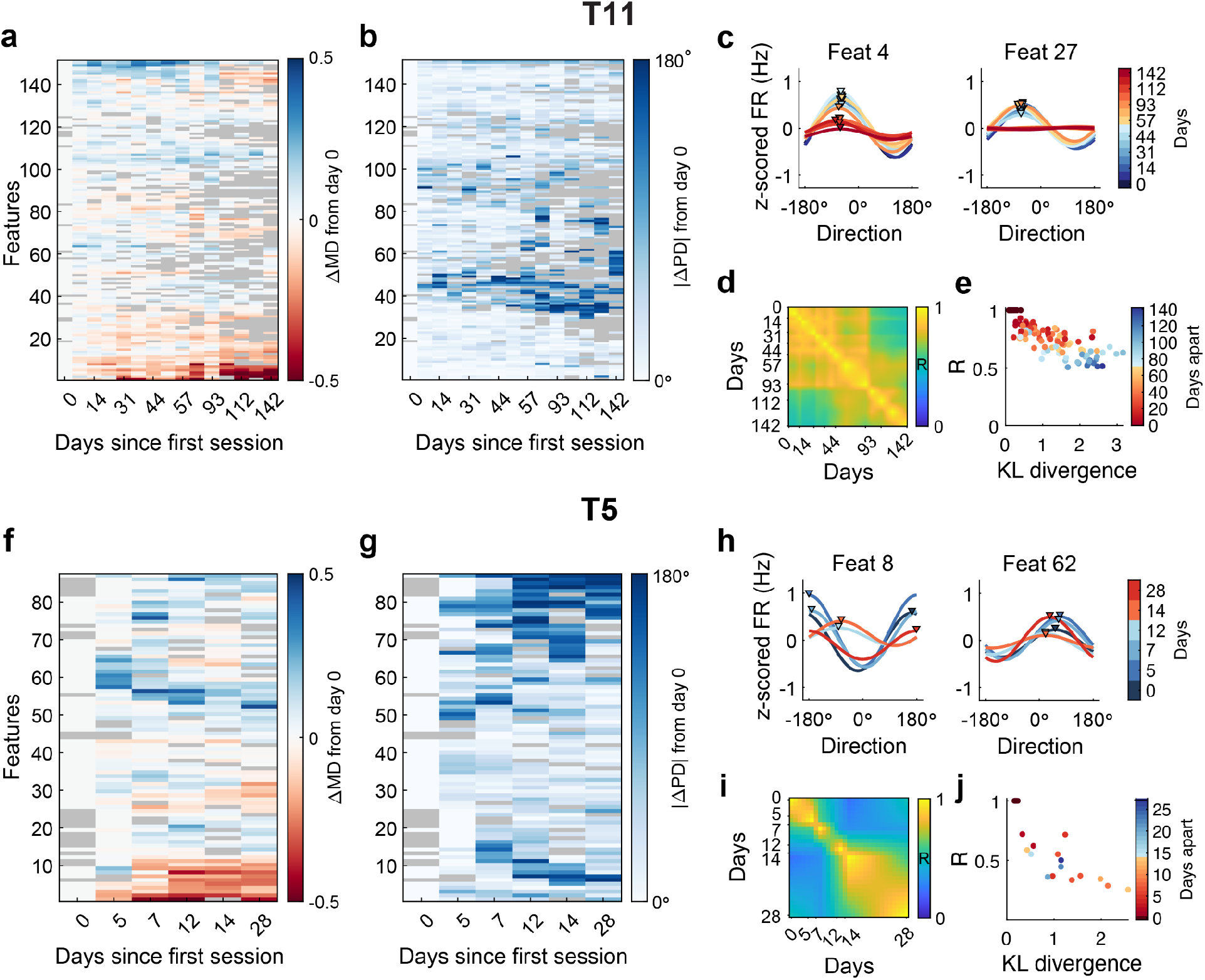
Changes in feature tunings across session correlates with KLD. **(a)** Changes in preferred directions (PD) and **(b)** modulation depth (MD) of significantly tuned features used in the decoder (relative to the tuning of the first day for which the feature was significantly tuned) for T11. Features were ordered by hierarchical clustering to visualize groups of features with similar tuning behavior changes (see **Methods**). Gray color indicates features that were not significantly tuned in that session. Triangle markers correspond to the features presented in (c). **(c)** Fitted cosine tuning curves for sample units across days for T11 illustrating changes in channel dropout and MD, respectively (color curves); Triangle markers denote PDs for sessions with significant tuning. **(d)** Tuning similarity across days represented by interpolated Pearson correlations between pairs of tuning maps (see **Methods**). **(e)** Mean KLD of neural distributions between sessions negatively correlates with the tuning similarity (Pearson *r* = -0.812, p < 10^−30^, see **Methods**). Each dot corresponds to a pair of sessions with the color indicating the number of days apart. **(f)** Changes in cosine tuning PD and **(g)** MD for significantly tuned features used in the decoder for T5. **(h)** Fitted cosine tuning curves for sample units across days of T5 illustrating changes in MD and PD, respectively. **(i)** T5 tuning similarity across session days. **(j)** Mean KLD of neural distributions between days strongly anti-correlates with the tuning similarity (Pearson *r* = -0.776, *p* < 10^−4^).

Similar to T11, gradual changes in T5’s tuning properties were also observed (**Fig. 3f-g**). Some features illustrated changes in either MD or PD, or both (**Fig 3h**). 71 out of 85 tuned features exhibited significant change in both MD and PD in at least one session. The average change in PD across these features was larger on days where performance was worse (day 12 and 14: 69.9°±40.7°; day 5, 7, and 28: 58.1°±36.6°; *p* = 0.0346, Wilcoxon rank sum). Average absolute change in MD was not found to be significant (day 12 and 14: 0.147±0.099; day 5, 7, and 28: 0.139±0.093; *p* = 0.664, Wilcoxon rank sum).

To quantify the changes in encoding on a population level, we use tuning maps ^56,58^, defined by matrices of fitted tuning parameters of significantly tuned features on each session. Tuning similarity between days was assessed by calculating the correlation between the corresponding tuning maps (see **Methods)**. In general, nearby sessions in time with similar performance were more correlated. For T11, tuning maps among early sessions (up to day 93) with high performance were highly correlated, as well as among later sessions with low performance, but not across these two epochs (**Fig. 3d**). For T5, sessions that were closer together in time (along the diagonal) had higher correlation than those further apart (**Fig. 3i**). These results suggest that model drift (tuning changes) occurred across sessions. We were thus interested in determining how the MINDFUL score reflects changes in tuning similarity. Instead of fixing a reference distribution, pairwise KLD of neural features between sessions were calculated using a sliding window of 60 seconds updating every 10 seconds (see **Methods**). The KLD from the same session were averaged to get a mean of the neural distribution difference between pairs of sessions.

For T11, pairs of sessions closer in time had smaller distribution shifts, while pairs of sessions further from each other in time had larger distribution shifts in terms of KLD, which consequently strongly correlated with tuning map correlation (Pearson *r* = -0.812, p <10^−30^, **Fig. 3e**). For T5, the same trend was observed, except for the pairs of sessions which compared the first three sessions to the last session where cursor control had recovered (dots in blue shades). The correlation between mean KLD and tuning map correlation was also strong and significant (*r* = -0.776, *p* < 10^−4^, **Fig. 3j**). Together, these findings suggested that the MINDFUL score using KLD captures day-to-day changes in directional tuning, even though the metric can be calculated without information reflecting target position or movement intentions.

### The MINDFUL score captures low-dimensional neural latent space drifts

We further investigated how this method relates to the changes in the low-dimensional neural latent space using demixed principal component analysis ^59^. The top two direction-dependent principal components (PCs) on neural population from decoder day 0 were calculated to compare changes across sessions by projecting the neural population from subsequent sessions in this PC space (see **Methods**). The average neural trajectories per target directions became less distinct as the session dates progressed, reflecting changes in the underlying population activity over time consistent with the decline in task performance (see **Fig. 4)**. The amount of direction-related neural activity in each session was quantified by the variance accounted for (VAF) by the top two direction-dependent components on the subspace of day 0. For T11, the VAF was initially 50.0% on day 0 and remained above 20% on days in which clear separation of target trajectories were observed. As the decoder performance declined, the VAF dropped to 4.4% on decoder day 142 (**Fig. 4a**). This change in neural representation in low-dimensional space is strongly and significantly correlated to the mean KLD per session (Pearson *r* = -0.892, *p* < 10^−5^). For T5, VAF was initially 42.2% on day 0, and subsequently dropped to 2.9% and recovered to 11.3% on the last session (**Fig. 4b**). There was also a strong and significant correlation between the top 2 VAF and mean KLD (Pearson *r* = -0.858, p = 0.0029).

**Fig. 4.**
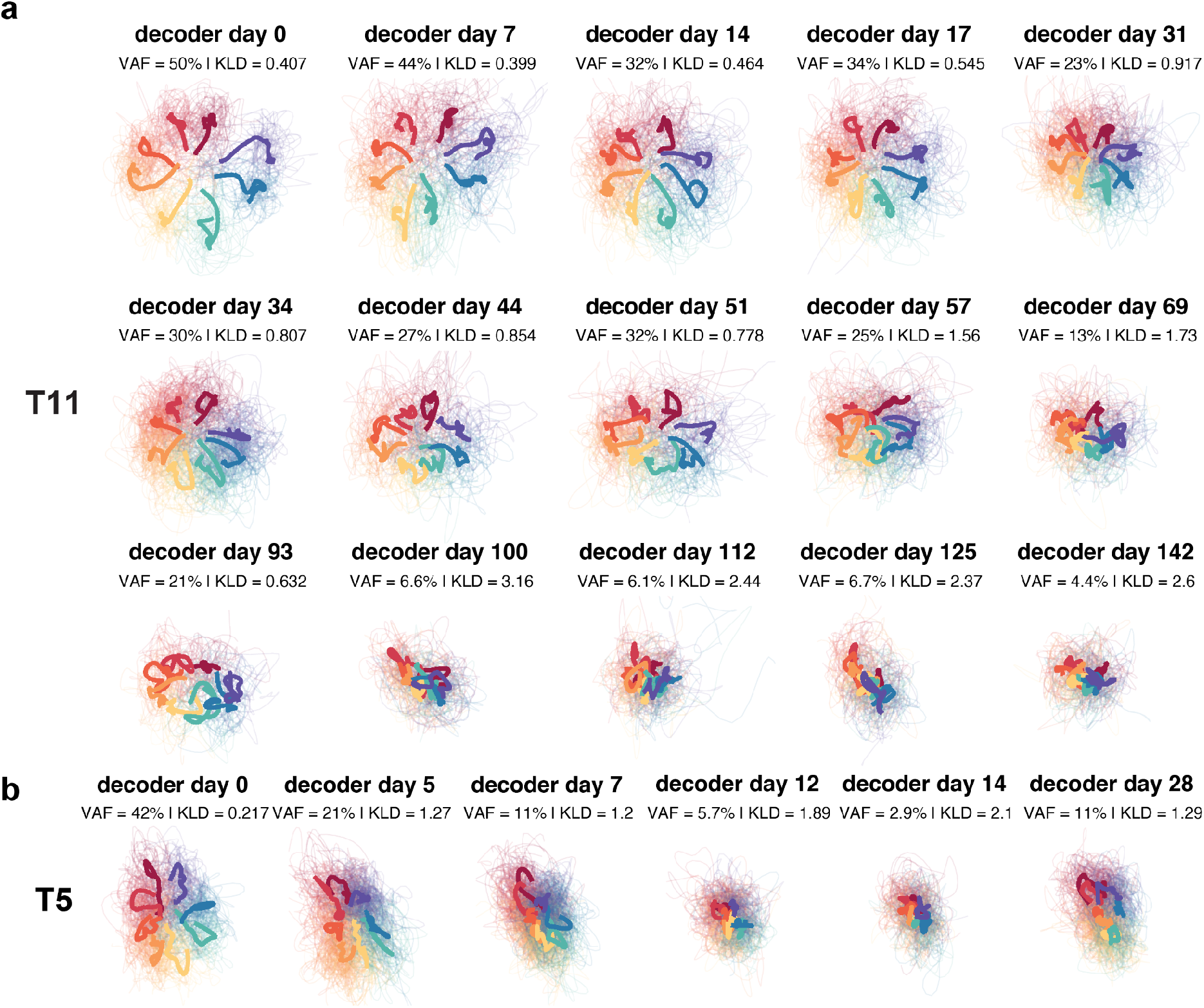
Instability reflected in neural latent space. **(a)** Projection of neural features of subsequent sessions onto the top two task-dependent PCs latent space of neural features on decoder day 0 using dPCA. Bold solid lines are trial averages per goal directions. Different colors correspond to the goal direction. **(b)**Projection of neural features for T5. For comparison simplicity, the random-target task was visualized and colored by discretizing goal directions of each trial into eight even movement directions as in a center-out-and-back task.

In addition to correlating with model drift in neural data, MINDFUL was found to detect large momentary deviations in the signal, likely attributable to device-related reasons such as signal transmission errors ^5^. The sharp spikes in both KLD and median AE in T11 neural data (**Fig. 2)** correspond to time steps during outlier trials (see **Supplementary Fig. 4**). Outlier trials were defined as having more than a 5% drop of wireless neural data packets or large “neural” responses greater than 8 standard deviations from the mean. Furthermore, when excluding these trials, the MINDFUL score was still highly correlated to the median AE (see **Supplementary Fig. 5**). This suggests that our method is capable of tracking both long-term neural instability, as well as short timescale technical-related variability. Although instantaneous events are less relevant for decoder recalibration, a method to capture these events may prove useful in other iBCI troubleshooting with both the current and future fully implanted systems.

### Selecting reference and window length further optimizes correlation

We explored the role of sub-selecting time bins with different AE ranges as the reference in the **MINDFUL** pipeline. When limiting the reference to collection of time bins with low AE only (0-4°) as shown in **Fig 1c, 2**, there are strong correlations between the KLD of derived neural features 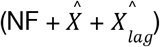 to AE. It was higher than when using all time steps of any cursor control quality for both participants (see **Fig. 5a**). This also held true when taking other combinations of derived neural features into calculating KLD (**Supplementary Table 1)**. In addition, sub-selecting instances with high AE (50°-100°, 100°-180°) as the reference distribution reduced the KLD-AE correlation for both participants, especially in T5.

**Fig. 5.**
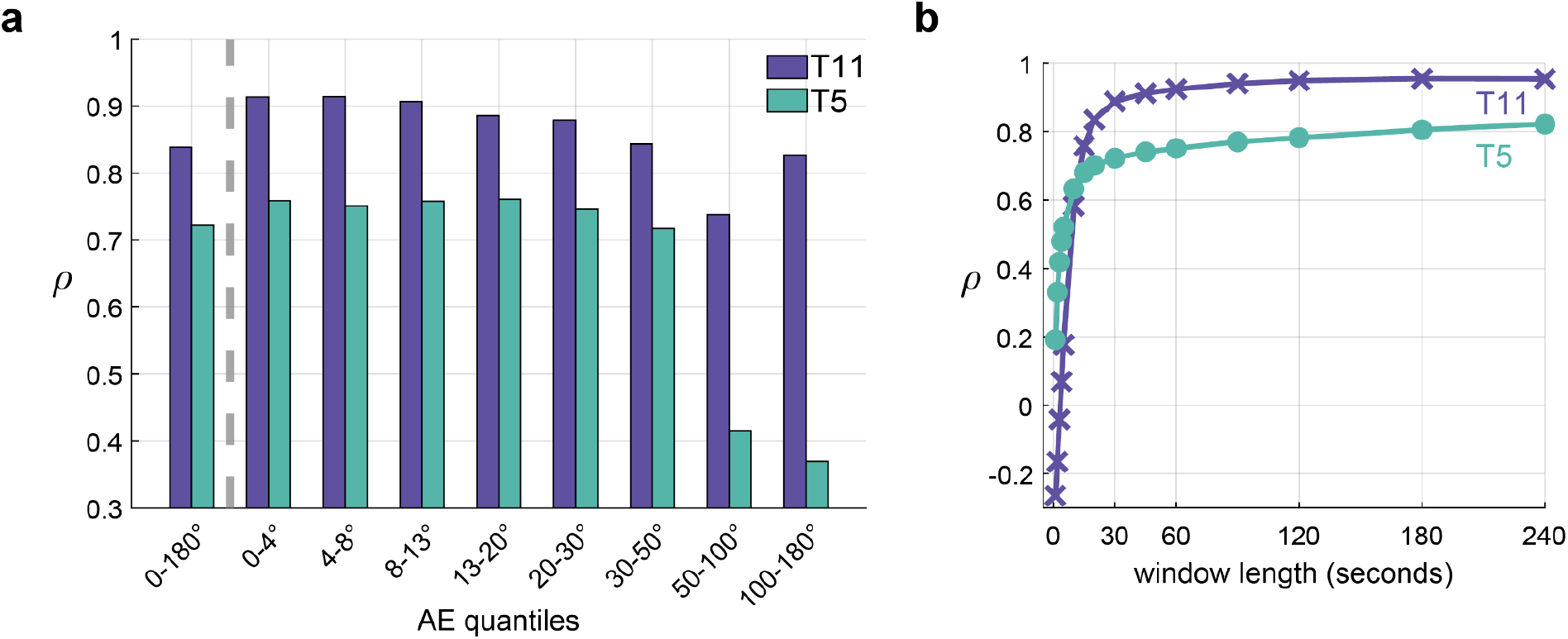
Sub-selecting instances of low AE for reference and using longer window length improve MINDFUL correlation to AE. **(a)** Spearman correlation coefficients of KLD of NF, 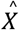, and 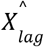, to AE when sub-selecting time steps with different quantiles of AE as the reference. Each quantile approximately contains an even number of observations. **(b)** Spearman correlation coefficients of KLD to AE when using different window lengths to estimate distributions.

Another important consideration for improving the MINDFUL correlation to performance is the duration of neural data required to obtain a reasonable representation of neural distribution. Since neural activity recorded in the precentral gyrus modulates with the direction of intended movement, too small of a window length may reflect task-dependent differences in reaching to different directions, rather than a persistent model drift that affects cursor control regardless of trial direction. A longer window length can avoid this issue when directional distribution differences are averaged out over a longer period. However, the KLD would be smoothed out if window length is too long, resulting in a larger delay to detect the need to update the decoder. To balance between accuracy and efficiency for online implementation, the optimal duration was empirically determined to be at least 60 seconds (**Fig. 5b**), where KLD-AE correlation began to plateau and longer windows did not offer a higher correlation. Using a 60-second window to estimate neural distributions provides a sufficiently large number of samples to average out directional-dependent differences due to variations in trial-to-trial movement directions.

## Discussion

Apparent model drifts during chronic iBCI use – resulting from changes in the information encoded in neural ensembles, changes in the recorded neural elements themselves, or changes in the recording devices – are one of the major challenges for developing decoders that will provide stable, accurate cursor control for long-term use by people with paralysis. Existing solutions to mitigate more substantial model drifts have limitations. Explicit recalibration comes at the expense of interrupting the user in the midst of iBCI use to collect additional data for training. While self-supervised recalibration doesn’t require the user to perform daily repetition of a task, it relies on stable online performance for effective pseudo-labeling. Currently, one cannot predict accurately the moment at which the decoder may fail to sustain performance and thus require a supervised recalibration. Similarly, robust algorithms reduce the need for frequent retraining but may require a retraining of the model *de novo*. MINDFUL fills in the gap in the development of a better decoder recalibration strategy for practical everyday use, by identifying, quantifying, and monitoring the degree of neural recording instability that contributes to degradation of real-time decoder performance.

The MINDFUL score, which is based on measuring the KL divergence (KLD) between distributions of neural features, reflects online performance without the need to incorporate knowledge of intended targets. In two participants, the MINDFUL score was strongly correlated to online angle error during iBCI cursor control across session days spanning up to four months (T11) or one month (T5). With the goal of translating this method to an online setting for the purpose of personal iBCI use, the MINDFUL score can reflect performance accurately for different cursor tasks examined in different participants (**Fig. 2**). Importantly, the MINDFUL score is consistent with tuning and latent space changes which can’t be directly measured without information about movement intention. This suggests that the MINDFUL score provides an intrinsic measure to track model drift affecting decoder performance during long-term iBCI use.

Our study confirmed the well-acknowledged observation that recording instabilities can impact online performance when the decoder cannot accommodate neural changes over long-term iBCI use. Model drift was quantified by tracking changes in tuning and latent space representations of neural population activity across sessions. MINDFUL was found to be highly correlated with both of these measures. It should be noted that our method did not track mean firing rate shifts which are known to correlate with declines in decoder performance ^19^. In our datasets, adaptive mean correction such as z-scoring or bias correction were applied to the neural features during online cursor control to combat this type of model drift (see **Methods**). Therefore, performance drops observed in this dataset were largely due to other types of model drift. The MINDFUL score, which measures changes in the distribution of z-scored neural data, aims to discover model drift such as changes in tuning or latent representation which may necessitate a decoder recalibration to restore control. Furthermore, since MINDFUL was designed to be applied during online iBCI control where threshold crossings were primarily used as neural features ^60–63^, we chose to investigate the functional stability in the thresholded neural activity’ tuning properties and latent neural representation rather than neuronal stability of discriminated single units processed via spike sorting techniques ^18,31^.

Another plausible contributor to the observed changes in neural distributions may be participants’ compensatory neuro-motor strategies in response to suboptimal cursor control. Non-human primates that encounter artificial perturbations in a previously learned BCI motor control task elicit new neural patterns with learning ^54,64–66^. Neural activity during closed-loop, online control contexts are also different from open-loop, offline control which has added real-time visual feedback ^67,68^. In this closed-loop iBCI study, when participants experienced a directional bias in cursor kinematics, it is plausible that the participants might compensate for decoding errors with different strategies such as moving against the bias (eliciting a larger magnitude of velocity), temporarily pausing attempting movement control, or moving towards the bias in hope to reset the bias (when automatic bias correction is applied). These alternative strategies are valid responses in attempting to improve control, but they may result in a larger change in distribution of neural features, amplifying the original model drift when intention context remains consistent. This highlights one of the challenges when studying model drift during closed-loop control when the ground-truth intended movement cannot be observed independently of the decoded outputs.

Selecting low AE time steps for reference was found to provide a higher correlation between the KLD and AE (**Fig. 5a**). Using low AE as reference helps to identify the model drift where the neural-kinematics conditional distribution during decoder training has changed from that during testing when the decoder was applied online. The training data of the decoder typically represents periods of relatively high performance: For T11, the LSTM decoder was trained on selected historical data with angle error < 45°, while T5’s decoder was initially trained on open loop blocks then immediately updated with a closed-loop block ^27,51^. As neural representations shift from the training distribution, the decoder is more likely to produce a subpar performance with a higher error rate. Therefore, when sub-selecting only high performance data as the baseline, future neural shifts can be more accurately reflected by the KLD. However, there exists subject variability and ambiguity regarding precisely how much data are needed for reference. For example, for T5, using the first two sessions as reference resulted in a slightly higher Pearson correlation than just the first session alone (**Supplementary Fig. 6**). Nevertheless, assuming that a newly trained decoder returns decent initial performance, our findings suggest that data from high-performance time steps from the initial session where the decoder was first applied would be an appropriate choice for the reference.

In this study, the MINDFUL score based on neural activity alone reflects performance in cases where fixed decoders were used online. But it remains unclear how it can be applied to other types of adaptive or robust decoders that aim to stabilize decoding by periodically realigning neural data to the initial session. In such circumstances, if MINDFUL is calculated before data alignment, it may not directly correlate to performance as adaptive alignment may keep AE low even when neural representations are changing. However, MINDFUL could still be useful in several ways. First, MINDFUL may be applied on transformed neural data after manifold alignment methods. Even if alignment approaches will result in a reduction in KLD (KLD is a common choice of loss function), MINDFUL can measure the remaining differences. Second, if the features used by MINDFUL are the outputs from the adapted decoder, rather than non-adapted features such as PCA components, then MINDFUL might continue to correlate to online performance despite decoder adaptation. Lastly, instead of periodically aligning (or recalibrating), MINDFUL can be used to signal the need for recalibration when model drift is detected. Future experiments with other closed-loop adaptive decoders will be required to test these approaches.

We proposed a statistical method to detect model drifts when fixed decoders were used during consecutive days-to-months of iBCI cursor control by two people with tetraplegia. This is crucial towards the goal of clinical translation of iBCI systems for practical everyday use, as it requires stable and reliable decoders to maintain high performance despite drifts in neural representations over time. MINDFUL was shown to be able to track model drifts based on the intrinsic properties of neural features and decoder outputs, which correlates to long-term changes in decoder performance, without needing to be aware of the movement intention. This approach is well-suited for future online adaptive iBCI systems aiming to provide continuous long-term control in a practical, personal setting outside of standardized research sessions, where it is not possible to directly track intended movements. For instance, MINDFUL and related methods could be used to trigger either a user-engaged or background update as the decoder begins to degrade. However, there are additional several considerations of applying the MINDFUL score online. First, during personal iBCI use, the movement directions could be less symmetric and more sparsely distributed than the cursor tasks used in this study. A longer time window or careful time bin selection may be needed to reduce the directional-dependent differences. Second, the threshold of KLD to determine when to recalibrate can be user specific. Lastly, the abovementioned large noise instances which can be easily detected by MINDFUL should not trigger a recalibration, as it does not imply a change in the neural-kinematic relationship estimated by the decoder. The frequency or pattern of these events could, however, be valuable in further iBCI development.

## Methods

### Human participants

The Institutional Review Boards of Mass General Brigham/Massachusetts General Hospital, Brown University, Providence VA Medical Center, and Stanford University granted permission for this study. Intracortical neural signals were recorded from participant T11, a 37-year-old right-handed male with a C4 AIS-B spinal cord injury (SCI) that occurred approximately 11 years prior to study enrollment, and T5, a 65-year-old right-handed male, with a C4 AIS-C SCI that occurred approximately 9 years prior to study enrollment. Both are enrolled in the BrainGate2 pilot clinical trial (NCT00912041), permitted under an Investigational Device Exemption (IDE) by the US Food and Drug Administration (Investigational Device Exemption #G090003; CAUTION: Investigational device. Limited by Federal law to investigational use). All research sessions were performed at the participant’s place of residence.

### Intracortical neural recordings and neural features

Each participant had two 96-channel microelectrode arrays (Blackrock Neurotech, Salt Lake City, UT) placed in the dominant (left) hand/arm knob area of the precentral gyrus ^2^. T11’s intracortical neural signals were recorded via a wireless broadband iBCI system ^69^ while T5’s recorded was acquired via the cabled iBCI system. All research sessions were performed at the participant’s place of residence. The average signal across the array per electrode was subtracted with a common average reference filter to reduce common mode noise. Neural features were extracted from the neural recording in 20 ms non-overlapping bins. For real-time decoding and offline analysis, multi-unit threshold-crossing spike rates (RMS < -3.5) per electrode were used for T5, and two types of features: spike rates (RMS < -3.5) and power in the spike-band (250 – 5000 Hz) per electrode were used for T11. Across the 15 sessions, 34 of the 1840 trials were labeled as outlier trials, which was defined as having more than a 5% drop of wireless neural data packets or large “neural” responses greater than 8 standard deviations from the mean. No outlier trials were identified in T5’s sessions.

### BCI behavioral task

To assess decoder performance for cursor trajectories, T11 performed a closed-loop 2D point-and-click center-out-and-back task for each of 15 sessions on separate days that spanned 4 months (trial days 658-800). For each trial, T11 was prompted to attempt hand or finger movements to continuously move the neural cursor from the center target to one of the eight pseudo-randomly selected peripheral targets and to then attempt a hand gesture (right index finger down) to click on the target. T11 was encouraged to maintain the same set of motor imagery for all sessions presented in this study. Upon target selection, in the next trial T11 was asked to move the cursor back to the center target. A trial is successful when the cursor is inside the target and a click action is decoded. Otherwise, a trial is considered failed after a 10-second timeout. Each session consists of two 5-mins task blocks, except for trial day 751 with only one block. The cumulative task time of all sessions is 145 mins, with a total of 1840 trials. Neural features, cursor position, target position, and decoder velocity outputs were logged.

T5 performed a closed-loop 2D random target selection task with a fixed-size target appearing in random locations on the screen. T5 attempted to move the cursor over the target and dwell on it for a consecutive 500ms period before the 10-second timeout to complete the trial. Audio feedback was provided right after the end of a trial to indicate trial success. A new random target is immediately presented with no delay. This task was repeated for 6 sessions on separate days that spanned 28 days (trial days 2121-2149). Each session consists of two to four 4-mins closed-loop blocks, which provide 84 mins of 1200 total trials across all sessions. Training blocks for calibrating the decoder on trial day 2121 were not included in the test data.

### Angle error

The instantaneous angle error is defined as the absolute angle difference between the inferred intended directional vector (cursor position to target position) and the decoded velocity vector, 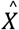 (best = 0°; max = 180°). In each 60-second interval to estimate the neural distribution for calculating the KLD, the median of the angle error time bins during the same interval is computed. Median is chosen over mean because AE in our datasets for both participants is not uniformly distributed between 0° and 180° (skew towards lower AE).

### Closed-loop neural decodings

Decoders in this study were previously described in ^27,51^. Briefly, for T11, an LSTM decoder was used to infer movement intentions from neural recordings. An LSTM is a variant of recurrent neural network (RNN) with improved capability for long-term temporal dependencies ^70^. Previous studies described the advantage of using a RNN for neural decoding over linear methods such as the Kalman filter ^20,71–73^. The LSTM decoder was trained and validated on closed-loop point-and-click cursor tasks from the 18 most recent sessions of T11, spanning 70 days from trial day 576 to 646. Of these sessions, only task blocks with a median angle error less than 45° were included, which yielded a total of 331 minutes or 8441 trials of training data. Input neural features were passed directly to the RNN layer whose outputs went to three densely connected activation functions, decoding the x- and y-velocity and the distance to target. During online control, each neural feature was adaptively z-scored using the mean and variance from a 3-minute rolling average window.

Clicks were decoded with a linear discriminant analysis (LDA) followed by a hidden Markov model. The LDA calculates a subspace that maximally discriminates between a click and a movement state. Coefficients were estimated with a regularization term of 0.001. Emission means and covariances used the empirical mean and covariance from the training data. The selected z-scored neural features were smoothed with a 100ms boxcar window before projected onto the LDA space. The estimated class probabilities were normalized using the SoftMax function, then smoothed with a 400ms boxcar window. A click is returned when the click probability was above a threshold of 0.98.

For T5, non-overlapping 20ms-binned extracted features were fed through a linear regression model trained to predict the cursor-to-target distance. An initial decoder was trained based on T5’s neural activity while he engaged in an open loop block on day 0 (trial day 2121). This decoder was then used to drive closed-loop control in a subsequent block. The final decoder parameters were then updated based on the first closed-loop block, and they were fixed for later closed-loop blocks and future sessions. The raw decoded velocity *v*_*t*_ was exponentially smoothed with the running velocity average 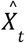 Via 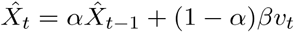, where α is the smoothing factor and Β is the gain parameter. Smoothing and gain were manually adjusted during the first session and fixed on subsequent days.

To accomodate for session-to-session variability in recordings, we applied per channel z-scoring at every time bin for T11 and a bias correction for T5. For T11, mean and variance were initialized from the previous block and adaptively update them using a 3-min rolling window. Neural features were decoded into cursor velocities by a real-time LSTM decoder. For T5, a bias correction was applied to mitigate mean shifts in the decoded output by subtracting a running estimate of the decoder bias from the velocity outputs (with an adaption rate of 0.3) ^11^. Bias correction was first initialized from the previous blocks. The intercept term in the decoder is then updated to the negative resulting bias vector (obtained by pushing the mean firing rate vector through the decoder weights).

### Derived neural features

MINDFUL can be based on any collection of neural features. In this paper we experiment with three different types of features. The first is extracted neural features as described above (384 dimensions for T11 and 192 dimensions for T5). Individual neural features were z-scored per channel using a 3-min rolling window as implemented during online iBCI control for T11. The same procedure is applied to T5 despite a bias correction approach was applied during online control to offset means drifts. The second is based on principal components analysis (PCA) of the extracted neural features (after z-scoring). The recorded neural features were projected onto the PCA subspace defined by the top M principal components (PCs) of a reference dataset that we call the *PCA-reference data*; see below. The third is based on the output of the decoder (which can be viewed as a type of neural feature), 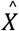. We also consider 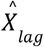, the previous time bin (20ms earlier) of 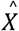.

### Kullback-Leibler divergence (KLD)

The MINDFUL score is based on comparing two datasets of neural features, defined in our case by the neural feature vectors from two collections of time bins. We use the derived neural features as described above. The first collection of time bins defines the reference *data P*_1_ and the second defines the *comparison data P*_2_. Our choices for the reference and comparison time bins are described below. We first compute the sample mean (column) vectors μ_1_ and μ_2_ of the neural feature vectors in the reference data *P*_1_ and comparison data *P*_2_, respectively. These mean vectors are the same dimension k as the derived neural features. We similarly compute the k x k sample covariance matrices Σ_1_ and Σ_2_ in the two datasets. The MINDFUL score that we use is

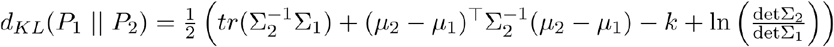

where tr and det denote the trace and determinant of a matrix, respectively, and ln is the natural logarithm. This formula is the Kullback-Leibler divergence (KLD) between two k-dimensional multivariate Gaussian distributions with respective mean vectors μ_1_ and μ_2_ and respective covariance matrices Σ_1_ and Σ_2_. Although it is motivated by a multivariate Gaussian model, its utility as a score for comparing two datasets does not rely on a Gaussian assumption. In developing MINDFUL, we experimented with other measures of statistical difference based only on means and covariances, such as the Jeffrey’s (symmetric KL), Bhattacharyya, and Wasserstein distances between multivariate Gaussians, and found qualitatively similar results (not shown). We selected KLD as the example for this paper, because it consistently gave the best results in many different scenarios and it is widely known.

### KLD grouped by angle error

For the results in **Fig. 1b**, the reference and comparison time bins are defined by the angle error (AE) of the time bin, regardless of movement intentio n and session date. The reference data consist of all time bins for which the AE < 4°. We used these same time bins for the PCA-reference data. The comparison data consist of time bins for which the AE is in a particular 4° interval. We used 45 different comparison datasets defined by the AE intervals [0,4°), [4°,8°), …, [172°,176°), [176°,180°] giving 45 different KLD scores, calculated as described above. These scores are plotted versus AE (using the middle of each AE interval for the AE value) in Fig. 1b, and these 45 pairs define the reported correlations for Fig. 1b. The derived neural features used are the top M=5 PCs.

### MINDFUL score to track model drifts during closed-loop iBCI control

MINDFUL quantifies the neural distribution shifts over time relative to a historical reference distribution. We use KLD as described above and experiment with different choices of derived neural features and reference and comparison time bins. The reference time bins are restricted to the first session when the decoder was first deployed for T11, and the first two sessions for T5 (the first session is shorter than the others; see **supplementary Fig. 6** for KLD using only the first session for reference). We additionally restricted the reference time bins to those with AE < 4° in **Fig. 1c, 2, 5b**. We varied this AE threshold for inclusion in the reference data in **Fig. 5a**. In all of these figures the PCA-reference data is the same as the reference data. **In Fig. 1c, 2, 5a**, the comparison data consists of all time bins in a 60-second interval. In **Fig. 5b**, we varied the length of the comparison data interval. In all of these figures the comparison data interval is shifted in increments of 1 second to investigate how the MINDFUL score varies over time. In **Fig. 1**, the derived neural features are the top M=5 PCs. In **Fig. 2a**, the derived features are the 2-dimensional decoder output 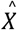. In Fig. 2b, the derived features are 4-dimensional and consist of 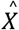 and the 2-dimensional decoder output from the previous time step 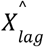. In **Fig. 2c, 5**, the derived features are 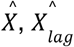, and the top M=5 PCs, for a total of 9 dimensions. **Supplementary table 1** shows results with additional choices of the derived features (no z-score, M=10, or no PCA).

### Cosine tuning

We fit cosine tuning curves to estimate the tuning properties per feature per session. Cosine tuning has been used to describe the relationship between the neuronal firing rate to movement directions, and it forms a basis of using a linear decoder for neural decoding ^57^. In a cosine model *y =b*_0_+*b*_1_ cos *θ* + *b*_2_ sin *θ, y*, the firing rate of a neuron, is regressed on θ, the movement direction. *b*_0,_*b*_1,_*b*_2_ are regression coefficients that can be estimated with least squares unbiased estimators. The model can also be expressed equivalently as *y = b*_0_+ *α* cos(*θ* −*θ*_0_), where 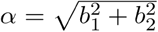 representing the modulation depth (MD) of the cosine curve. and *θ*_0_ representing the preferred direction (PD) where the largest firing rates are recorded. It is noted that since features in this study were z-scored, the bias term is closed to zero, and is therefore omitted in the calculation of modulation depth. Tuning parameters were estimated from 20ms-binned features from the first second after the go-cue of non-outlier trials (offset by 160 ms reaction time) to capture neural activity associated with reach initiations. Feature tunings are considered significant if the F-test on the regression model has a p-value < 0.05. See **Supplementary Fig. 7** for the estimated tuning and the empirical firing rate of example features.

### Changes in MD and PD

For each feature on each day, change in MD was calculated by Δ *MD*=*MD*−*MD*_*ref*_, and change in PD was calculated by |Δ *PD*|=min(360°−*PD*_*ref*_|, |*PD* − *PD*_*ref*_),where *MD*_*ref*_ and *PD*_*ref*_ refer to the tuning of the first day for which the feature was significantly tuned. The CircStat toolbox was used ^74^. Only features that have significant directional tuning for more than half of all sessions were considered in the tuning map described below. (For all features including the non-significantly tuned, see **Supplementary Fig. 8**). Significance of tuning change was assessed by bootstrapping samples of PD (or MD) to obtain a distribution of ΔMD and ΔPD ^33,35^. If the 95% confidence interval for the difference distribution does not contain 0, then we reject the null hypothesis at the 5% significance level ^35^. To visualize the patterns of tuning changes, the features were ordered by their tuning parameters using hierarchical clustering on Matlab. Euclidean distance was used to estimate the similarity of standardized ΔMD and ΔPD from all sessions between two features, and ward linkage was applied to arrange the order of the clusters. The same ordering was used in the heatmap of ΔMD and ΔPD for the same participant.

### Changes in tuning map

We quantified changes in directional tuning on a population level by comparing tuning maps over recorded sessions. A tuning map on each session is a 3 x *N* matrix comprises the fitted tuning curve parameters, *b*_0,_*b*_1,_*b*_2_, of *N* number of features with significant tuning on that day. Pairwise Pearson correlations of maps was performed to assess the similarity of tuning across sessions. In a pair of daily tuning maps, only features that were significant on both maps were considered to calculate the correlation. This pairwise correlation was plotted in a heatmap which was interpolated to account for the irregular number of days apart between sessions.

### Mean KL between sessions

To estimate the average neural distribution difference between pairs of sessions, a mean KLD between distributions on two given sessions was calculated (see **Fig. 3e, 3j**). The 9-dimensional derived neural features are the same as **Fig. 2c, 5**, namely, the top M=5 PCs, 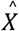, and 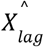.

The PCA-reference data is the same as **Fig. 1c, 2, and 5**, namely, time bins from the first session (T11) or first two sessions (T5) that have AE < 4°. The reference and comparison data are each 60-second intervals updated every 10 seconds. If there are M such intervals in session *i* and N in session *j*, then the mean KLD between the sessions is the average KLD of all (*M* × *N*) pairs of intervals from sessions *i* and *j*. Outlier trials were excluded from the intervals. For visualizing the complete pairwise comparison matrix containing the KLD of all pairs of intervals from all sessions, see **Supplementary Fig. 9**.

### Latent space dPCA projection

We applied demixed principal component analysis (dPCA) ^59^ to population neural activity from the initial decoder day for T11 and T5 respectively. For subsequent trial days, we projected neural data onto the top two task-relevant neural dimensions dPCA space of day 0. PCs were computed from all features of the first second after the go-cue of non-outlier trials (offset by 160 ms reaction time). Neural features were smoothed using a gaussian kernel with standard deviation of 50 ms. T5 random-target task trials were discretized into eight movement directions in order to show comparable results to T11’s center-out-and-back task with eight peripheral targets. We further quantified the amount of task-related neural activity in each session by comparing the variance accounted for (VAF) by the top two task-related neural components from the first session. VAF was computed by

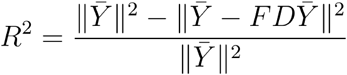

where 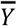 is trial-average neural features across conditions on a subsequent day, and encoder components *F* and decoder components *D* estimated from the first session.

## Code Availability

The code for reproducing the figures will be made available on https://github.com/ewinapun/MINDFUL.

## Author contributions

We have included a graphical representation of author contributions below.

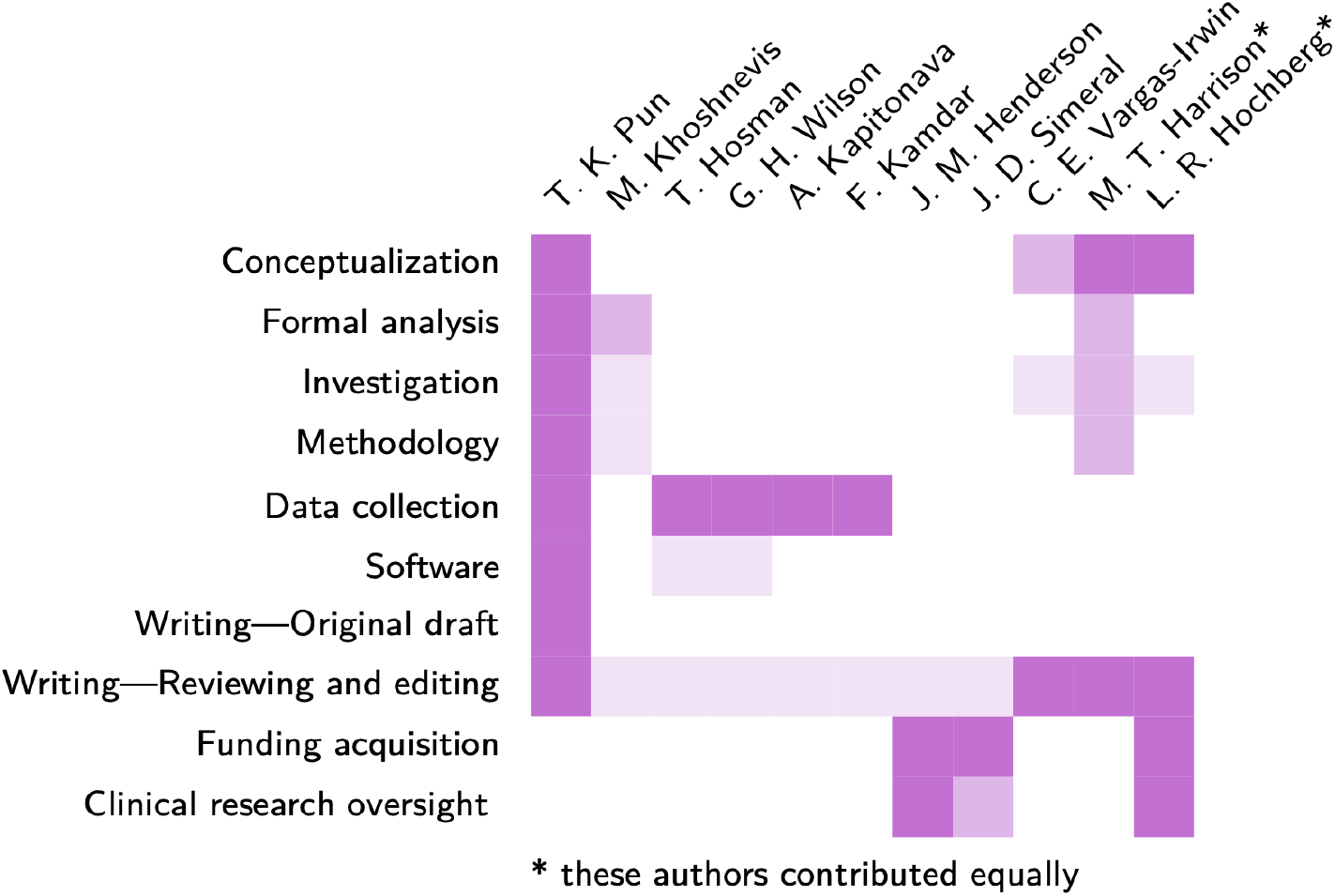

## Acknowledgement

The authors would like to thank participants T11, T5, their families and caregivers, Beth Travers, Dave Rosler, Maryam Masood, Ronnie Gross, Kristi Emerson, Sandrin Kosasih, Beverly Davis, and Kathy Tsou for their contributions to this research. We acknowledge Krishna Shenoy for his inspiration and effort toward creating an environment that spawned this work. Research was supported by the Office of Research and Development, Rehabilitation R&D Service, Dept of Veterans Affairs (N2864C, A2295R, A2827R, A3803R), NIH NIMH (T32MH115895), NIH-NINDS (UH2NS095548, U01NS098968), NIH-NIDCD (U01DC017844, R01DC014034), the Croucher Foundation, Larry and Pamela Garlick, Wu Tsai Neurosciences Institute, Howard Hughes Medical Institute (HHMI) at Stanford, and Simons Foundation Collaboration on the Global Brain 543045, Hong Seh and Vivian W. M. Lim endowed chair and HHMI Investigatorship at Stanford University.

## Supplementary materials

**Supplementary figure 1.**
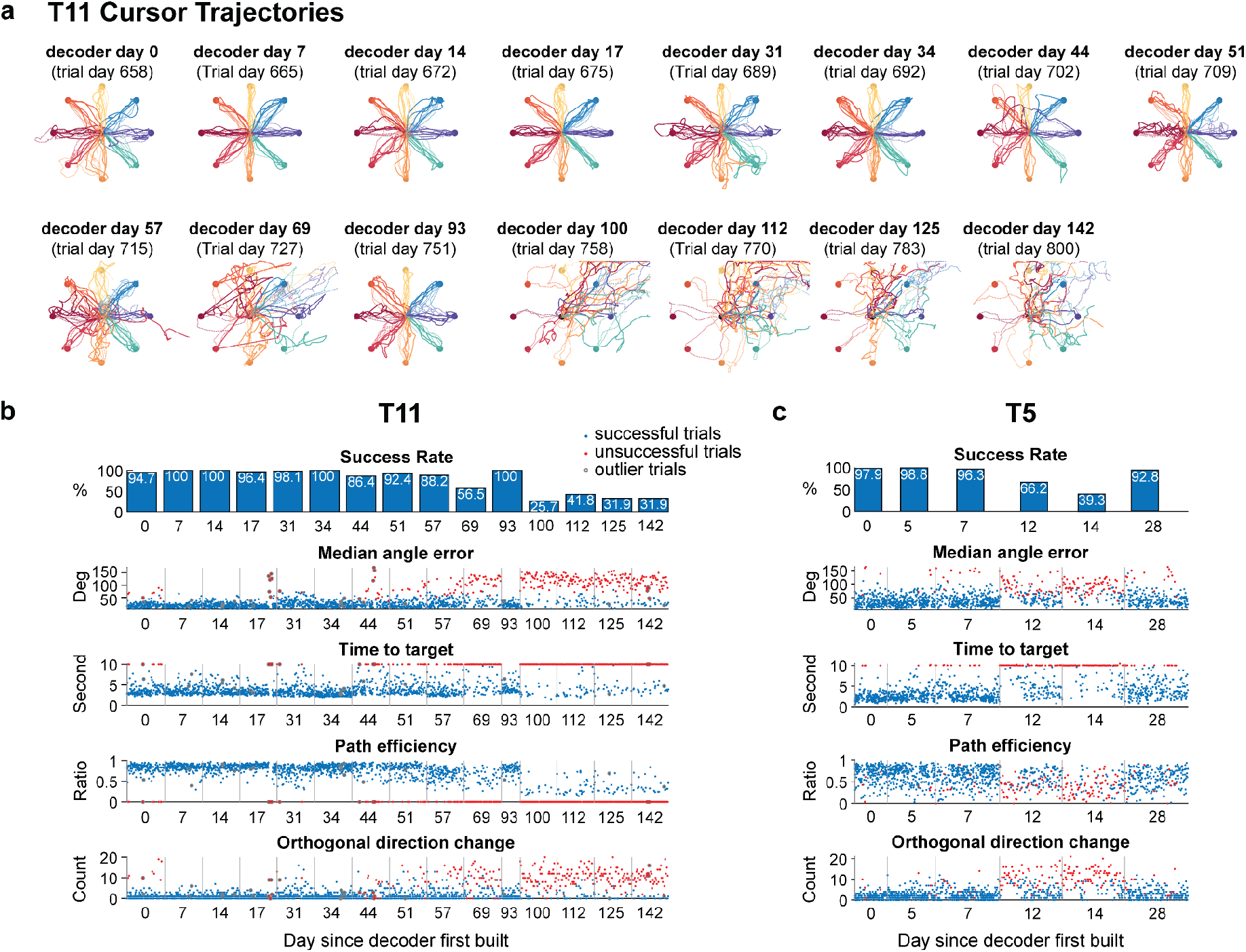
Cursor task performance using a fixed decoder. **(a)**Adapted from ^51^. Cursor trajectories in the first 5-min center-out task for selected trial days. Colors correspond to different target locations. A solid line indicates the cursor trajectory of a trial going from center to a peripheral target, and a dotted line indicates cursor trajectory of a trial from a peripheral target back to the center. Towards later sessions where the decoder began to degrade for T11, there was a consistent directional bias to the upper right corner. Trials that required the cursor to move to the bottom-left targets tended to fail. **(b)**Adapted from ^51^. Overall trial success rate per trial day, and other trial-to-trial performance metrics across trial days for T11. Each dot represents either a successful trial in blue or an unsuccessful trial in red where the cursor fails to reach the target before a 10-second timeout. Trials with gray circles are outlier trials with significant noise in the recordings. Cursor control is assessed using the following performance metrics: trial success rate, average trial success, angle error; time to target, time to reach to the target per trial; path efficiency, which is the ratio of the actual trajectory length to the Ideal straight-line path (best = 1) in a trial; and the number of orthogonal direction changes (ODC), where the cursor reversed away from or then back toward the target, which quantifies the path consistency towards the target (best = 0). The decoder achieved an average of 93.8% success rate in reaching and clicking the cued target in the first 11 sessions in the three months (trial day 658-751). But it subsequently degraded to 33.1% in later sessions (trial day 758-800). On average per trial, early sessions (658-751) demonstrated higher performance than later sessions (758-800) in terms of median AE (early:26.8°±22.6°; later: 88.4°±46.1°; *p* < 0.001; Wilcoxon rank sum), time to target (early: 4.02±1.98 seconds; later: 8.01±3.04 seconds; *p* < 0.001), path efficiency (early: 0.79±0.12; later: 0.38±0.17; *p* < 0.001; excluding unsuccessful trials) and orthogonal directional change (early: 1.82±2.82; later: 7.40±5.35; *p* < 0.001). **(c)**Same performance metrics across trial days as **(b)** for T5. There were no outlier trials that had significant noise events. Trial success rate in the first three sessions was 97.6% compared to 71.6% in later sessions. The first three sessions demonstrated higher average performance than the later three sessions in terms of AE (early: 39.6°±23.9°; later: 58.8°±31.7°; *p* < 0.001), time to target (early: 2.87±1.71 seconds; later: 5.90±3.12 seconds; *p* < 0.001), path efficiency (early: 0.66±0.20; later: 0.52±0.22; *p* < 0.001; excluding unsuccessful trials) and orthogonal directional change (early: 2.21±2.08; later: 5.59±4.65; *p* < 0.001).

**Supplementary figure 2.**
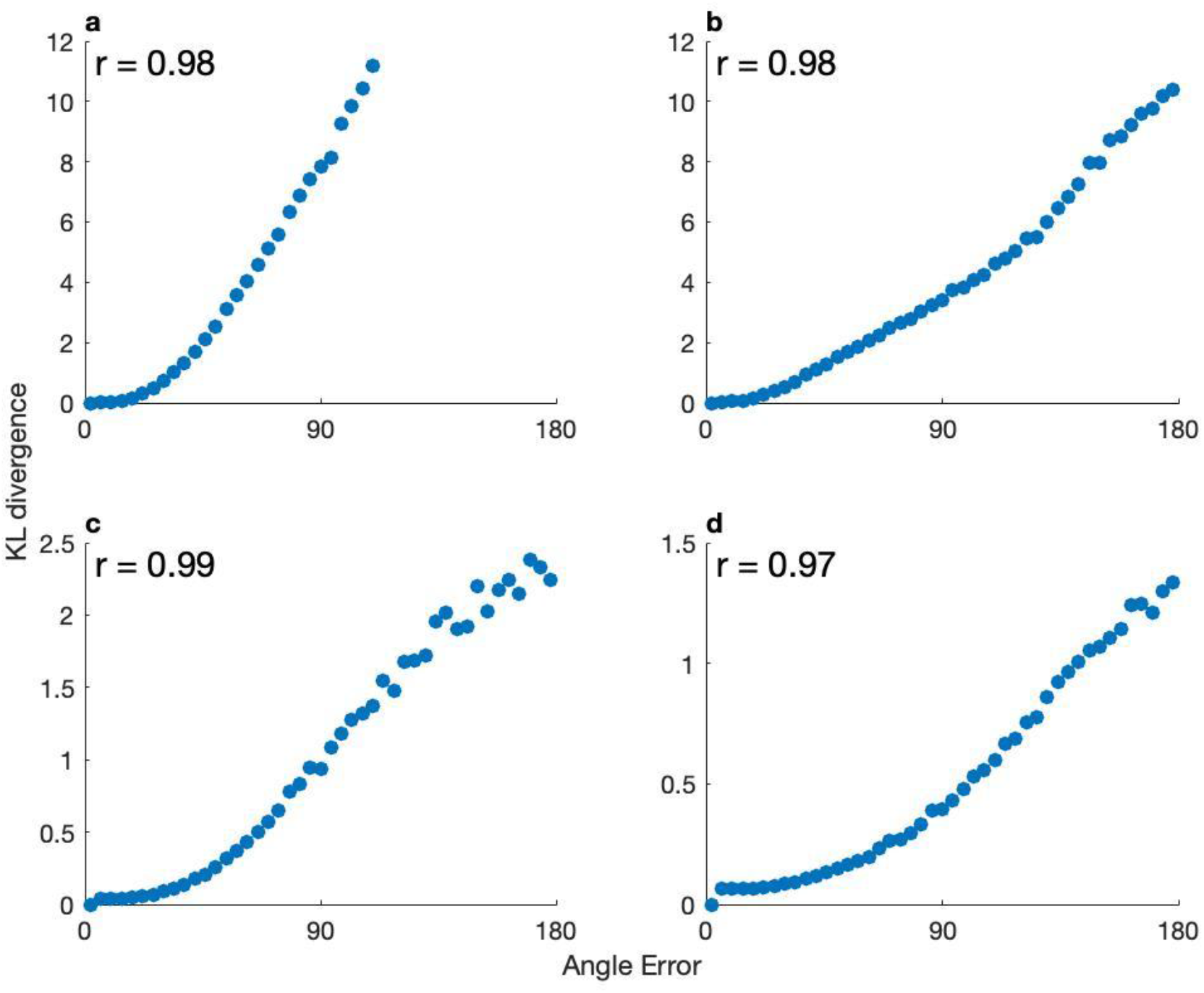
A linear relationship between KL divergence and angle error in simulated noise or model drift. *KL*(*p*(*X*|0<*AE*<4),*p*(*X*|θ−2<*AE θ*+2)) changes with angle error, (see **Supplementary methods**). **(a-b)** displayed case I where noise was applied without additional model drifts, and **(c-d)** displayed case II where added model drifts (tuning changes) were applied. In **(a), (c)**, we use a low model noise (*σ*^2^=4) (), while in **(b), (d)**, a higher model noise (*σ*^2^=25) is applied. Pearson correlation coefficient was shown in each panel. Both simulated noise and model drift result in a linear relationship between *KL*(*p*(*X*|*AE*≃0) and *p*(*X*|*AE*≃*θ*)) with *θ*. Additionally, a higher level of noise corresponds to increased angle error. The difference in coverage of angle errors is notable: **(c)** spans the entire possible range (from 0° to 180°), while panel **(a)** only extends up to 110°. This discrepancy potentially underscores the contribution of model drifts to the KL divergence, surpassing mere noise effects.

**Supplementary figure 3.**
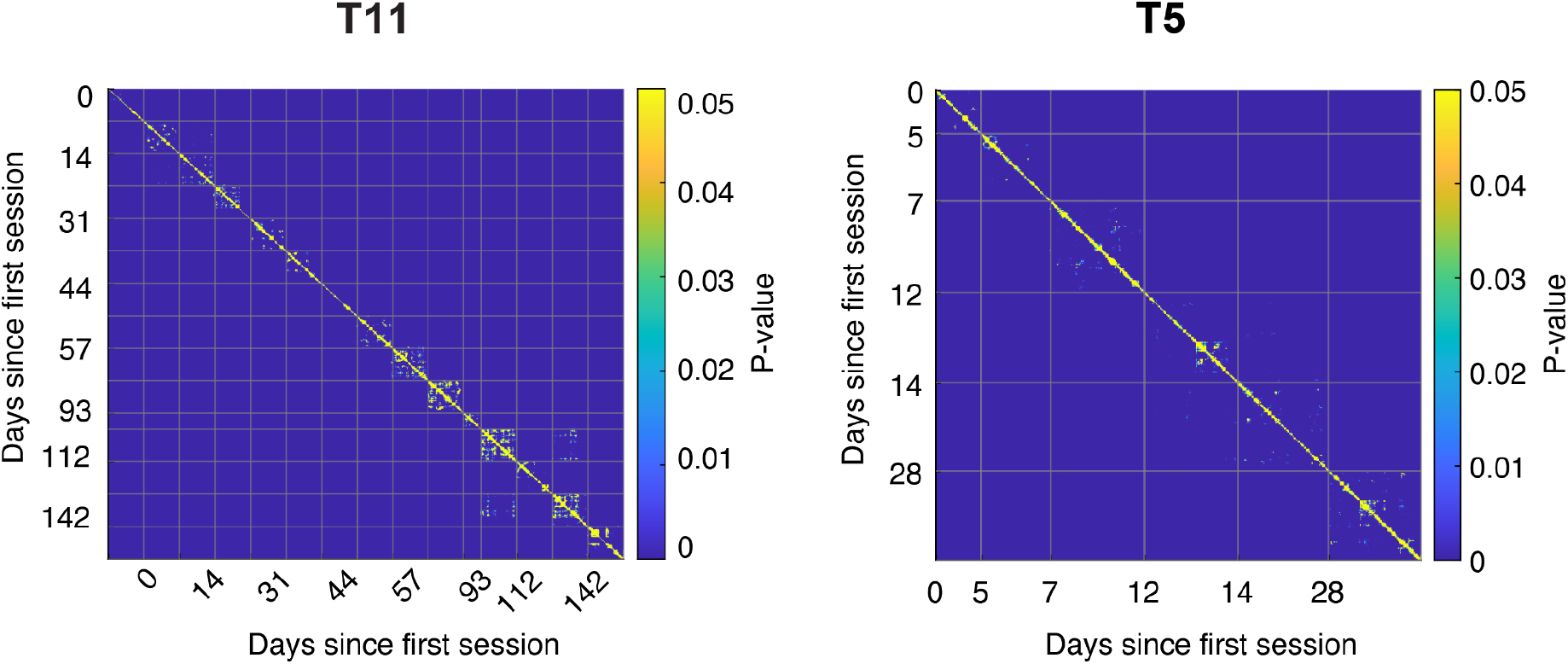
Multiple hypothesis testing with Bonferroni corrected p-value. Bonferroni corrected p-value between pairs of kinematics-neural data from time bins from sessions was computed, for T11 and T5 (see **Supplementary methods**). The compared data are each 60-second sliding window updated every 10 seconds. The data used to obtain the PCA transform are the same as **Fig. 1c, 2**, and **5**, namely, time bins from the first session (T11) or first two sessions (T5) that have AE < 4°. Neural features are projected to the top M=5 PCs. The hypothesis test suggests the presence of model drifts on almost every pair, except the neighbor ones along the diagonal, for both participants.

**Supplementary figure 4.**
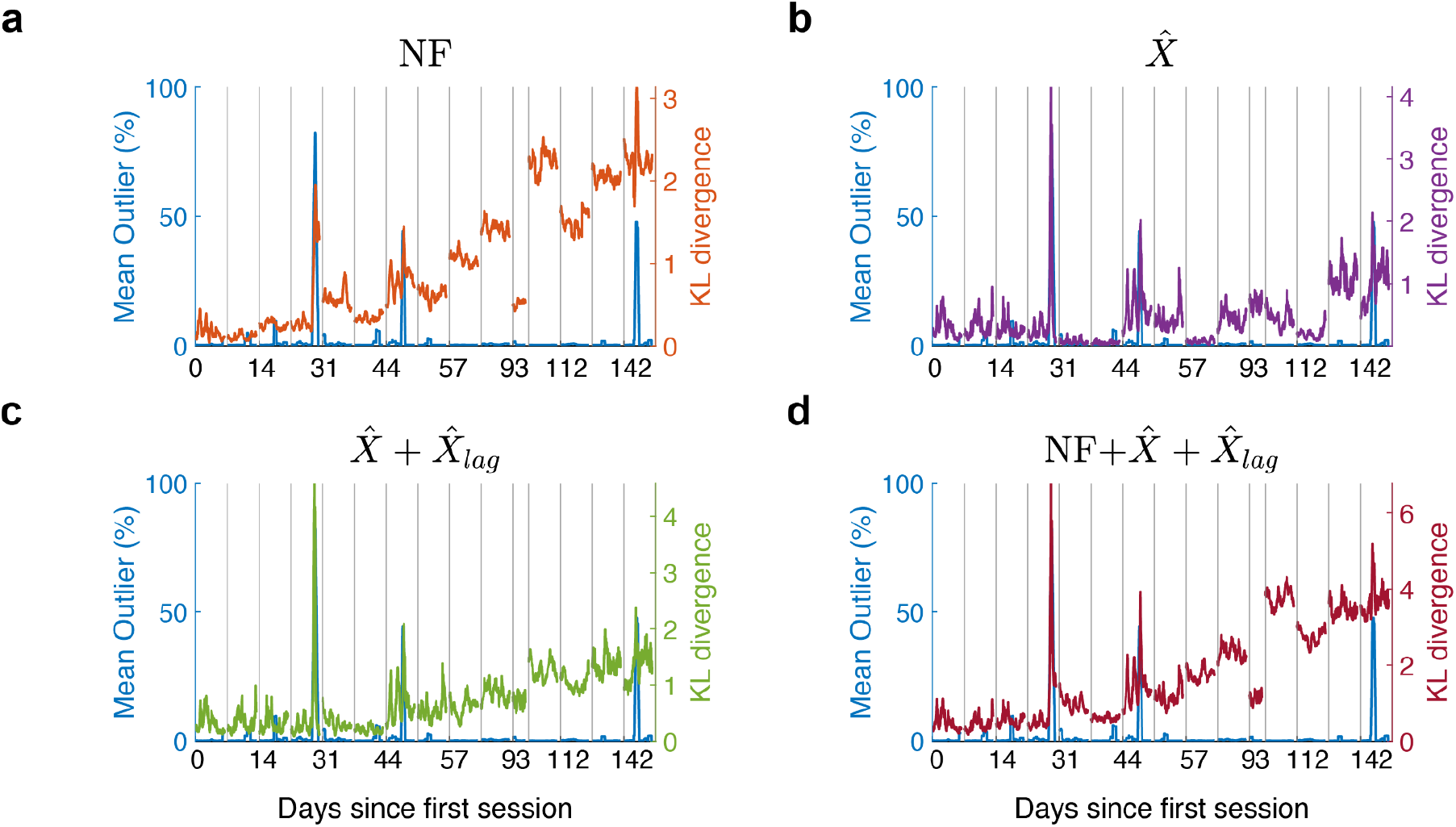
Sharp spikes in the KLD correspond to recording outliers, Participant T11. Mean outliers over each sliding window of 60 seconds were shown on the left y-axis. Outliers at every 20 ms time bin were defined by the maximum percentage of noise events across electrodes during a session. Noise events can be attributed to drop packets and bit flips from data transmission and large recording signal instances (>8 standard deviations from mean). Gray lines indicate the beginning of the session. **(a)** Across all sessions, large spikes were observed in the KLD on low-dimensional neural data (same as **Fig. 1a**) at the occurrence of three large abrupt momentary surges of outlier events (around end of day 17, middle of day 44 and 142). Large spikes in KLD using **(b)** decoded velocity, 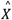, or **(c)** 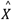and its lag,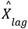. were presented for the first two outlier surges on day 17 and 44. **(d)** When KLD was computed on low-dimensional neural data along with decoded velocity and its lag, the large spikes on day 17 and 44 was prominent whereas the spike on day 142 is less prominent than **(a)**. This suggests that large noisy events were reflected in the recorded neural data, and can affect the decoded cursor movement but only momentarily. And KLD appeared to be sensitive to these large sudden deviations due to noise.

**Supplementary figure 5.**
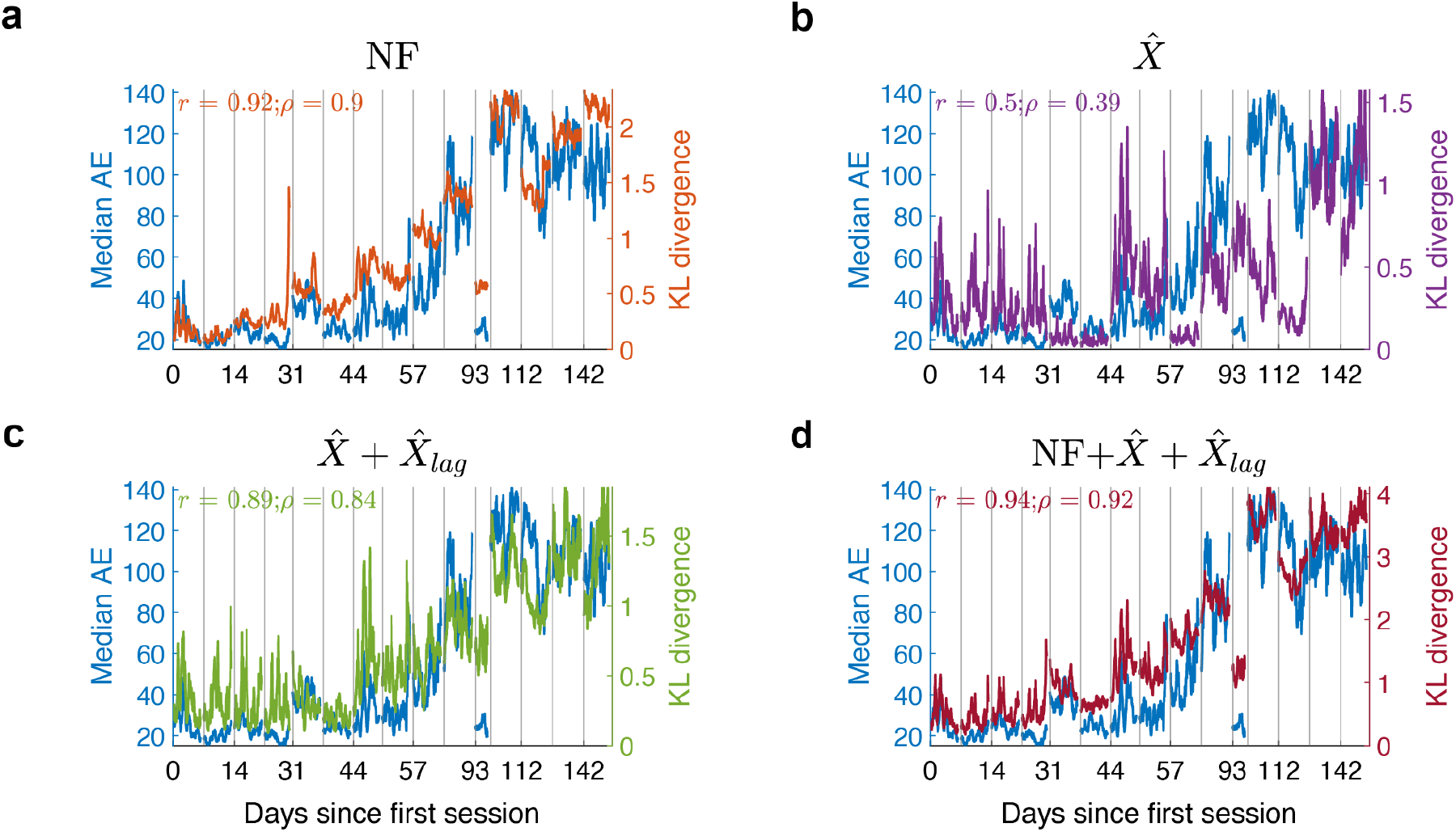
KLD excluding the outlier trials, Participant T11. When excluding trials with more than 5% mean outliers across the trial duration, KLD was also strongly correlated with the online median AE, and even higher than when outlier trials were included. This reflects that MINDFUL was still capable of monitoring long-term model drift that leads to degrading performance, in addition to detecting outlier events.

**Supplementary figure 6.**
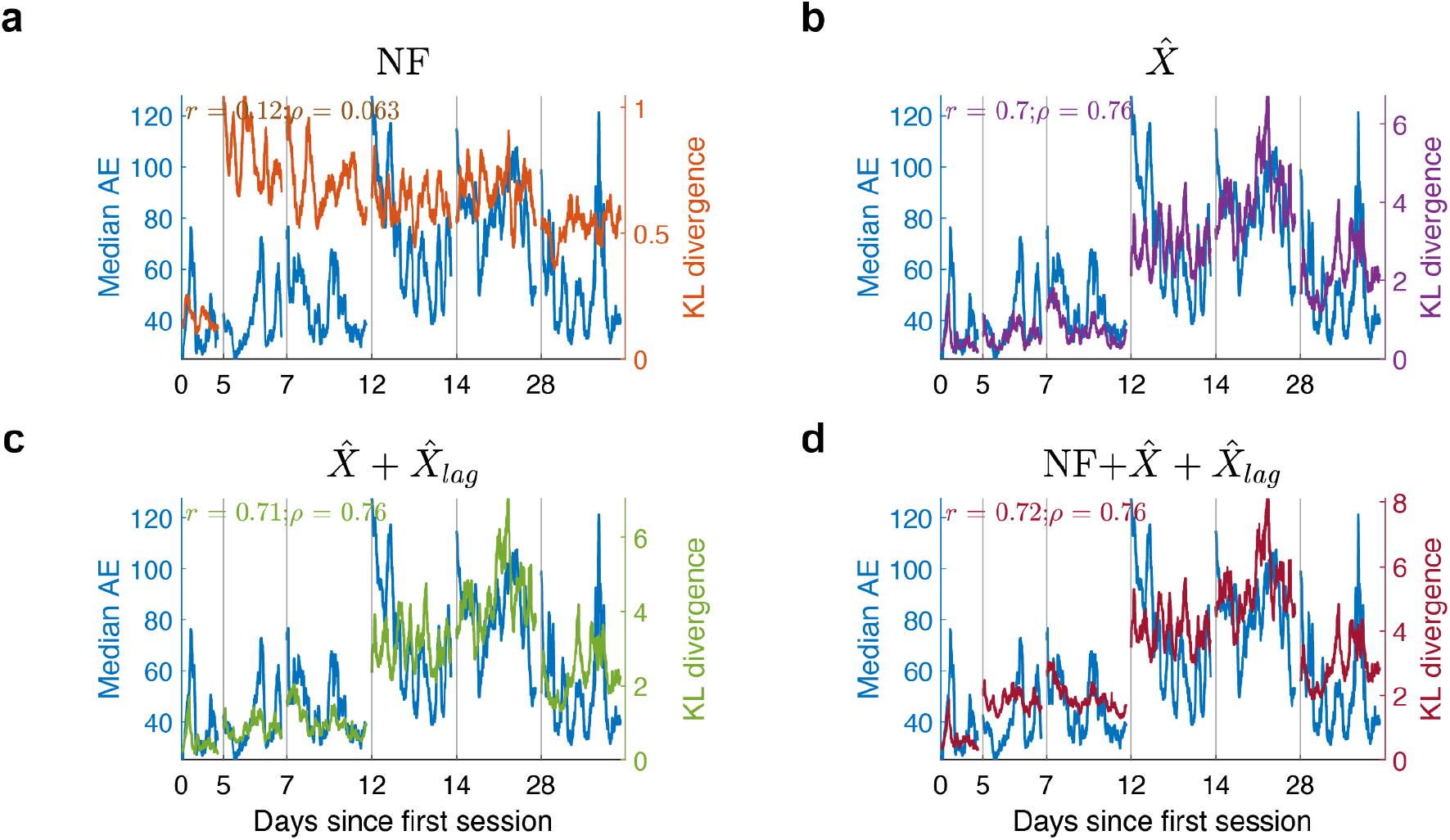
KLD using the first session as the reference, Participant T5. **(a)** KLD (right y-axis) overlaid onto median angle error (left y-axis in blue) across all recorded sessions T5. Distributions were estimated from low-dimensional neural data after PCA. Neural features on the first session were selected for the reference distribution. Only time steps when AE < 30° were sub-selected as the reference. Subsequent neural distributions and median AE were updated every 1 second over a 60-second sliding window. Pearson *r*, and Spearman rank correlation coefficients ρ, between KL and median AE were calculated. **(b)** KLD between distributions of just decoded directional vectors,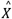, not neural data. **(c)** KLD between distributions of just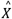 and 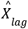. **(d)** KLD between distributions of a combination of low-dimensional neural data, decoded directional vectors,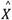, and 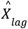.

**Supplementary figure 7.**
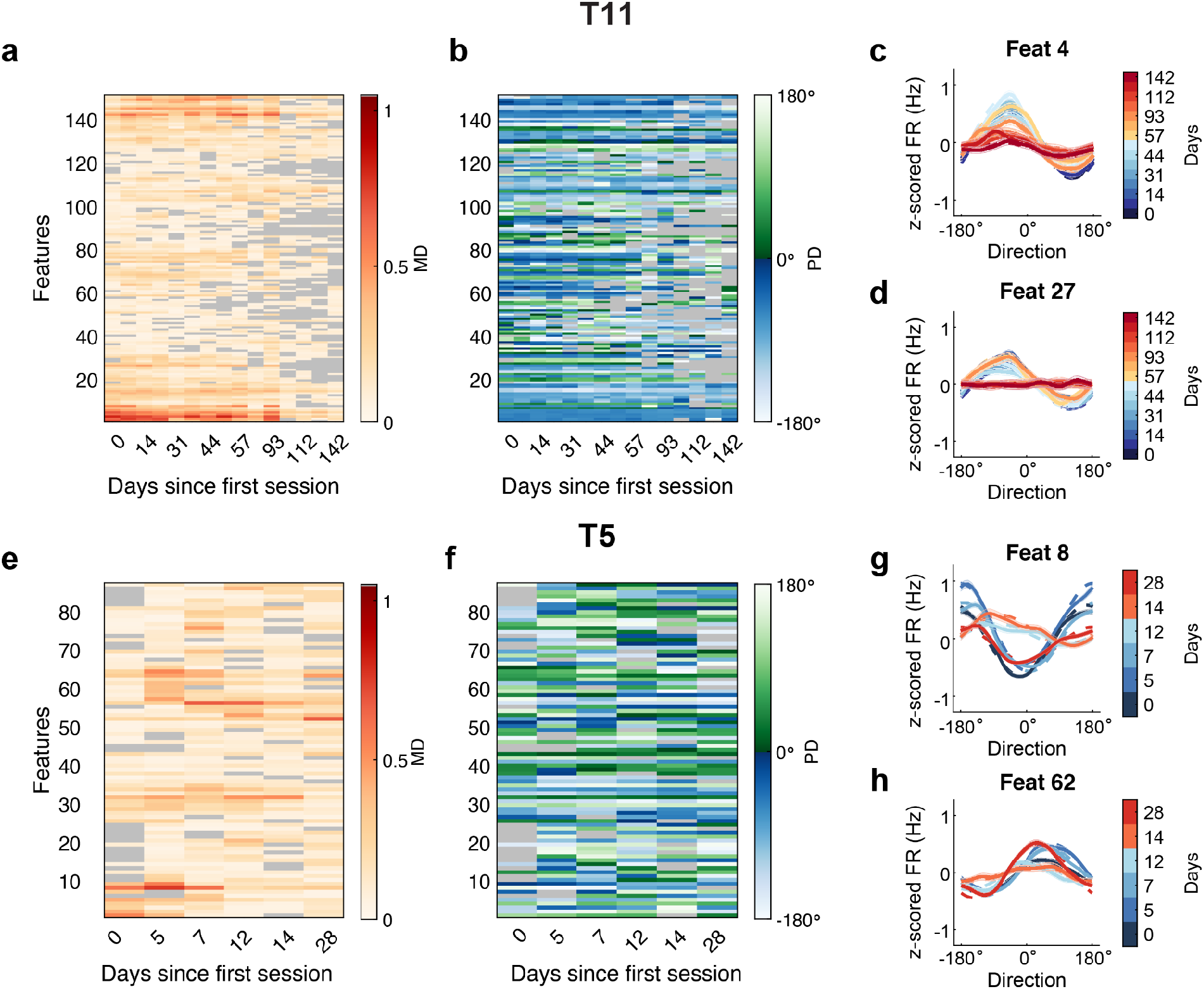
Modulation depth (MD) and preferred direction (PD) of individual features. **(a)** MD and **(b)** PD of significantly tuned features of T11. Features are sorted using the same hierarchical ordering as **Fig. 3(a)** and **3(b). (c-d)** Example neural model drift of two features across sessions of T11 along with empirical firing rate. Dotted lines correspond to cosine tuning models fit within each session as shown in figure 4d and 4e. Solid lines are empirical firing rates estimated using Nadaraya-Watson kernel regression, along with the 95% confidence intervals in shaded bands ^75^. Line colors denote the session day from light to dark progression. **(e)** MD and **(f)** PD of significantly tuned features of T5. **(g-h)** Example neural model drift of two features across sessions of T5 as shown in **Fig. 3(i) and 3(j)**.

**Supplementary figure 8.**
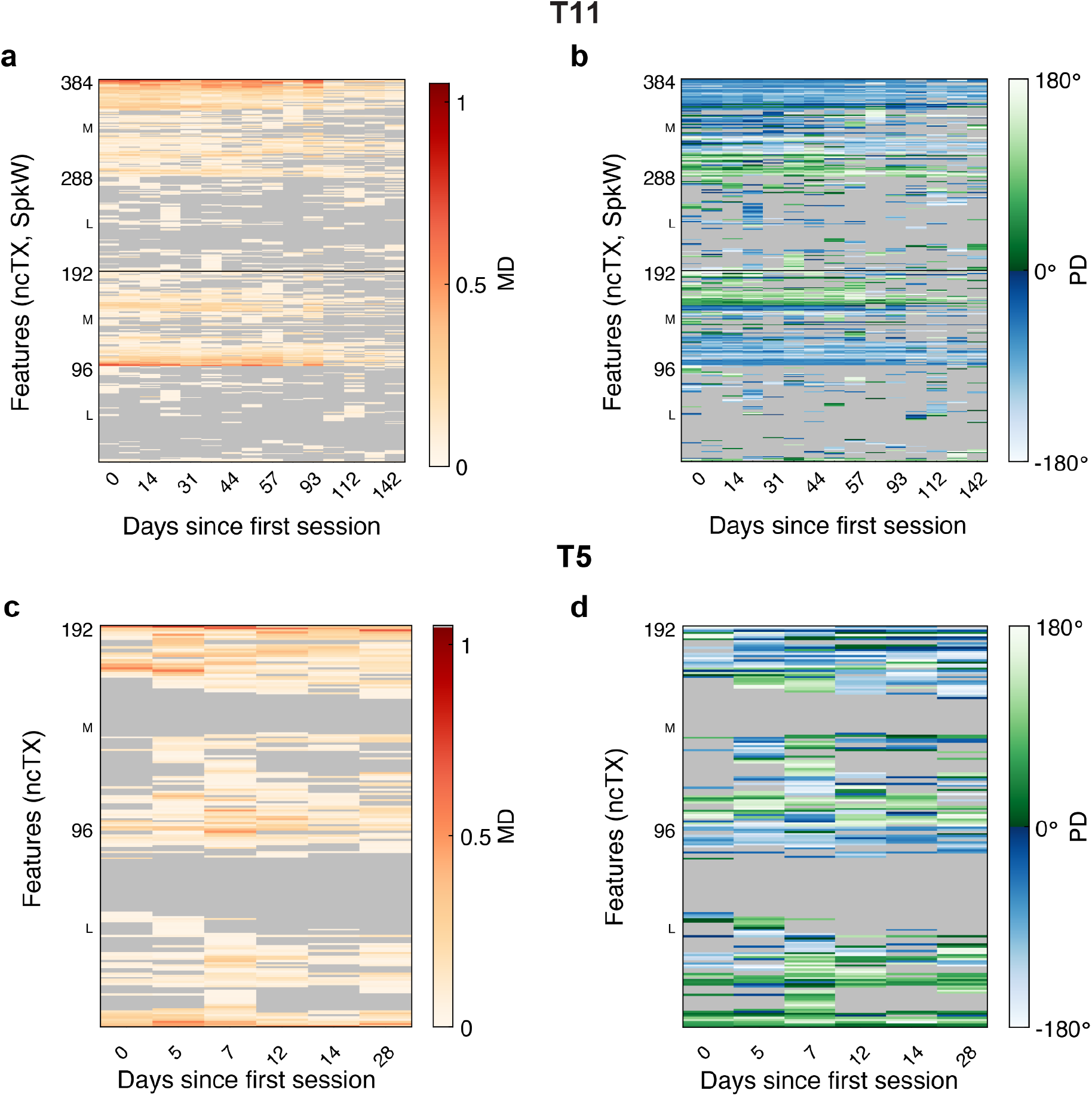
Modulation depth (MD) and preferred direction (PD) of all feature tunings. Tunings for all features used for online decoding were estimated. **(a)** MD and **(b)** PD of all features used in online decoding across all sessions for T11. Both tuning maps were ordered using the same hierarchical clustering methods. Hierarchical clustering was applied per array (L: lateral, M: medial) per feature type (ncTX: threshold crossings, SpkW: spike power). Gray out features were not significantly tuned (p > 0.05). **(c)** MD and **(d)** PD of all features sorted by hierarchical clustering per array for T5.

**Supplementary figure 9.**
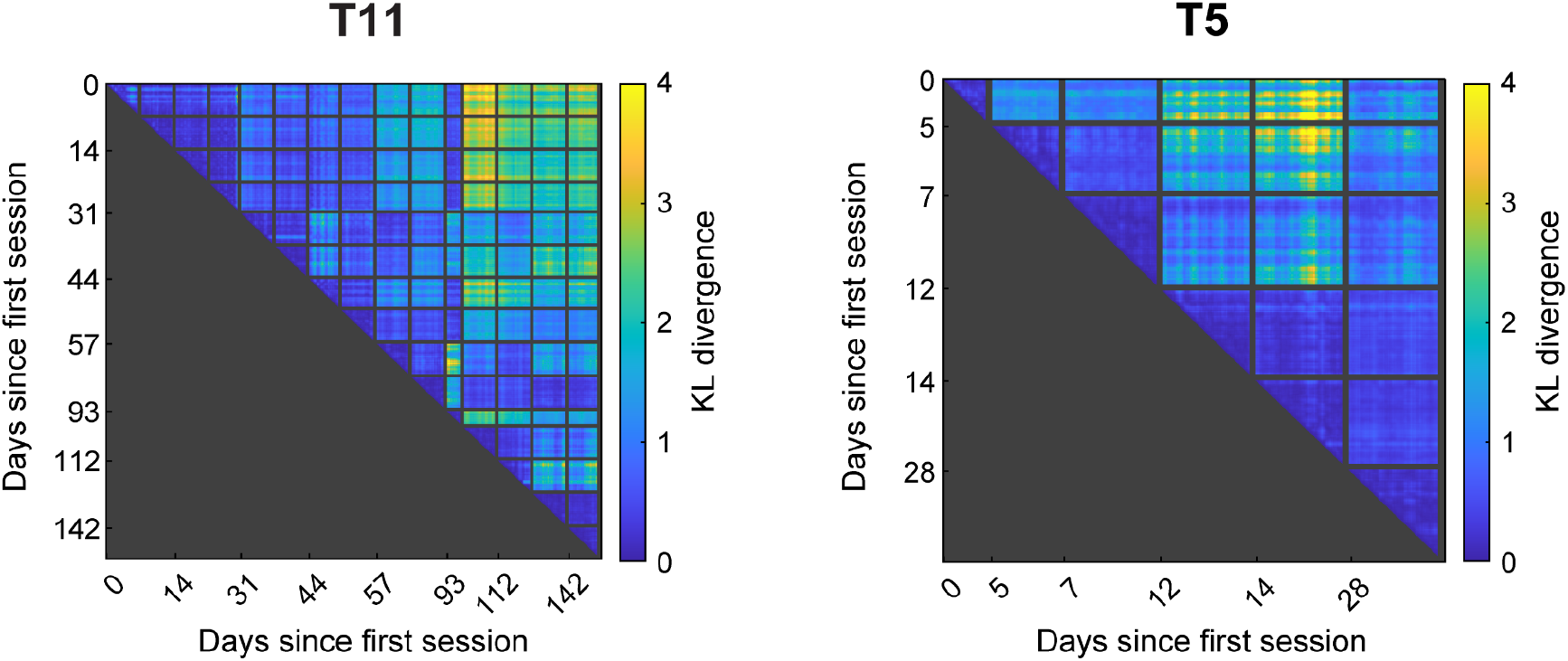
Pairwise KLD between sessions. For each day, distributions were estimated from low-dimensional neural features, in combination with decoded output, 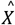and 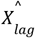, from overlapping time segments using a 60-second sliding window updated every 10 seconds. Outlier trials were excluded. Only the upper triangle matrix is shown as KLD is computed between distributions estimated from two time segments where the first segment precedes the second segment chronologically.

**Supp. Table 1.** Ablation analysis for MINDFUL. Higher dimension in neural features (NF): Using 10 PCs rather than 5 when estimating the mean and variance of the high-dimensional neural data did not make a difference in the correlation between the KLD and AE. No reference sub-selection: Without sub-selection of time steps with low AE (<4°) in the reference distribution, correlations decreased slightly for cases where NF were used as part of the input features to calculate the KLD. However, correlations were higher for T11, and similar for T5 when only using 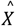 or 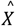+ 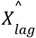 as the features. No z-score in NF: Without z-scoring neural data, mean shifts were more pronounced in measuring distribution difference using KLD (not shown). This resulted in low correlation to online performance since mean shifts were accounted for with adaptive z-scoring and bias removal for T11 and T5 respectively. No PCA in NF: When not applying PCA, there was little-to-no correlation in terms of Pearson’s coefficient for both T11 and T5 in cases where NF were used. This could be due to the fact that directly estimating the distribution of high dimensional data becomes difficult as the number of samples in a 60s sliding window is relatively limited when compared to the number of dimensions. It also substantially increases the computational power required.

|  | Fig. 2 |  | Higher dim.<br>in NF |  | No reference<br>sub-selection |  | No z-score<br>in NF |  | No PCA<br>in NF |  |
| --- | --- | --- | --- | --- | --- | --- | --- | --- | --- | --- |
| Reference<br>sub-selection | AE < 4° |  | AE < 4° |  | all timesteps |  | all timesteps |  | all timesteps |  |
| PCA: top PCs | 5 |  | 10 |  | 5 |  | 5 |  | - |  |
| T11 |  |  |  |  |  |  |  |  |  |  |
| Correlations | $r$ | $\rho$ | $r$ | $\rho$ | $r$ | $\rho$ | $r$ | $\rho$ | $r$ | $\rho$ |
| NF + $\hat{X} + \hat{X}_{lag}$ | 0.926 | 0.913 | 0.923 | 0.898 | 0.923 | 0.838 | 0.319 | 0.602 | 0.093 | 0.522 |
| NF | 0.91 | 0.9 | 0.905 | 0.877 | 0.887 | 0.774 | 0.325 | 0.596 | 0.09 | 0.52 |
| NF + $\hat{X}$ | 0.878 | 0.882 | 0.9 | 0.882 | 0.902 | 0.801 | 0.321 | 0.596 | 0.091 | 0.52 |
| $\hat{X}$ | 0.464 | 0.407 | 0.464 | 0.407 | 0.69 | 0.748 | 0.69 | 0.748 | 0.69 | 0.748 |
| $\hat{X} + \hat{X}_{lag}$ | 0.819 | 0.84 | 0.819 | 0.84 | 0.827 | 0.844 | 0.827 | 0.844 | 0.827 | 0.844 |
| T5 |  |  |  |  |  |  |  |  |  |  |
| correlations | $r$ | $\rho$ | $r$ | $\rho$ | $r$ | $\rho$ | $r$ | $\rho$ | $r$ | $\rho$ |
| NF + $\hat{X} + \hat{X}_{lag}$ | 0.719 | 0.759 | 0.72 | 0.758 | 0.706 | 0.722 | 0.387 | 0.375 | 0.025 | 0.152 |
| NF | 0.587 | 0.596 | 0.53 | 0.532 | 0.474 | 0.498 | 0.244 | 0.289 | 0.016 | 0.129 |
| NF + $\hat{X}$ | 0.715 | 0.768 | 0.716 | 0.765 | 0.717 | 0.775 | 0.391 | 0.377 | 0.019 | 0.188 |
| $\hat{X}$ | 0.704 | 0.763 | 0.704 | 0.763 | 0.708 | 0.78 | 0.708 | 0.78 | 0.708 | 0.78 |
| $\hat{X} + \hat{X}_{lag}$ | 0.702 | 0.765 | 0.702 | 0.765 | 0.702 | 0.749 | 0.702 | 0.749 | 0.702 | 0.749 |

### Supplemental methods

#### 1 Linear KLD-AE relationship using simulated models

The timescale of how a change persists matters in clinical settings. Because if the dominant cause of the linear relationship demonstrated in Fig 1b is intermittent noise, which occurs at sub-second timescale and does not contain any persistent changes, one cannot predict performance by combining time bins of neural activity in any way without knowing the performance in the first place. However, if the dominant cause comes from changes that persist over time, i.e. model drift, any contiguous group of time bins of neural data may define a collection where its statistical properties will reflect a shared source of error.

To study how each noise and model drifts contributes to KLD, consider the following a linear regression encoding model for neural data given kinematics,

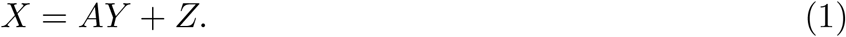

In this equation, *X* represents the matrix of neural features (size *d* × *n*), *Y* is the movement intentions (size *l × n*), and *A* denotes the regression coefficients (tuning) matrix (size *d* × *l*). Here, we used *d* = 50, and *l* = 2. We assume a multidimensional Gaussian noise *Z* in this model, which is independent of *Y* and follows a distribution of *N*_*d*_(**0**, *σ*^2^*I*).

For this linear model, under the assumption that *Y* ∼ *N* (**0**,*I*), the optimal decoder 𝔼[*Y* |*X*] is 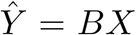, where *B* = *A*^*T*^ *(AA*^*T*^ *+ σ*^2^*I*)^-1^. We denote *AE* = angle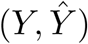 as the angle error of this estimation.

We generate synthetic data by considering two scenarios:

**Case (I)**: No added model drifts:

We simulate neural features using the encoding model in (1) for *n* = 10^4^ timesteps for 50 simulated sessions. Throughout these sessions, the parameters of the model *A* and *σ*^2^ remain constant. This process yields the matrix *X*, representing neural features of size *d* 50n. After decoding *X* using the optimal decoder *B = A*^*T*^ (*AA*^*T*^ *+ σ*^2^*I*)^-1^, we calculate the angle error for each time step across sessions.

Next, we estimated two conditional distributions:

1. The target distribution of neural features *X* conditioned on a specific angle error range *θ* - 2 < *AE* < *θ* + 2, and,
2. The reference conditional distribution of neural features *X* given good performance, where 0 < *AE* < 4. We compute the KLD for varying angle errors θ ∈ {2, 6, 10, …, 178}.

**Case (II)**: Model drifts with tuning changes is added:

The process remains akin to the one in case (I), except for a daily alteration in the parameter *A* of the encoding model. This parameter changes randomly each session day from the original tuning matrix *A* (used to build the decoder *B* = *A*^*T*^ (*AA*^*T*^ + σ^*2*^*I*)^-1^) to a newly chosen tuning matrix *A*^*^. This alteration is achieved using a linear combination of the original matrix and the new tuning matrix: *λA*+(1 -*λ*)*A*^*^, where *λ* is uniformly chosen at random from [0, 1]. Despite this daily change in the encoding model, the neural features *X* across all days is still decoded using the same decoder *B*, which was specifically optimized for *λ* = 0. (The other parameter of the encoding model (*σ*^2^) remains unchanged daily). Similar to case (I), the KLD between *p*(*X*|0 < *AE* < 4) and *p*(*X*|*θ* - 2 < *AE* < *θ* + 2) (both assumed Gaussian) for varying *θ* is calculated.

This procedure captures how the encoding model’s parameter changes affect the distribution of neural features, even when decoded using an optimal decoder designed for a stable parameter (*λ* = 0). The KLD is computed across different angle error ranges (*θ*) to assess the impact of these parameter variations. In contrast, in case (I), where there is no change in the encoding model parameters, the KLD is solely affected by the instantaneous noise present in the system.This distinction allows us to discern how variations in the encoding model parameters specifically contribute to the differences observed in the neural data distribution, separate from the impact of mere noise fluctuations.

#### 2 Hypothesis test for model drifts

As another method to detect model drifts in our probabilistic encoding model, we use a multiple hypothesis test described as below.

For two kinematics-neural pairs (*Y, X*) and 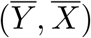, where *X* and 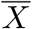 are *d*-dimensional neural features of sizes *d* × *T* and *d* × 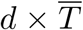 respectively, and *Y* and 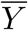 are movement intention matrices of size 2 × *T* and 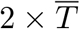, we assume the linear regression models below:

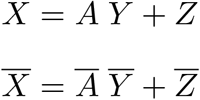

where *A* and 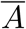 are the tuning matrices, and each column of 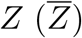 is independently distributed as 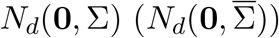.

The null hypothesis which we want to test is 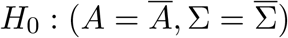. Under the assumption that *Z* and 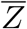are independent, we follow Toyoda and Ohtani(1985) to define two hypothesis tests for each dimension of neural features separately. In other words, for each dimension 1 ≤ *j* ≤ *d*, since 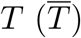 observations are assumed to be independent, the models above can be written as

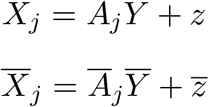

where 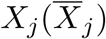 and 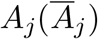 denote the *j*’s row of 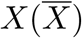 and 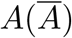, and 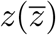is a 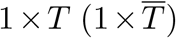 matrix of noise distributed as 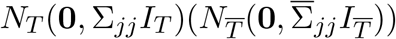. (Σ_*jj*_ is the (*j, j*) entry of Σ.) Now for each 1 ≤ *j* ≤ *d*, we use test statistics given in [1] to find 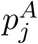 and 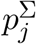, *p*-values associated to the individual tests 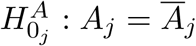 and 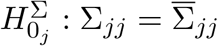 respectively in the following way:

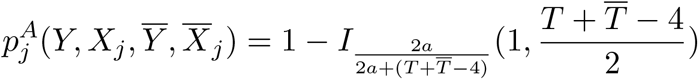

for

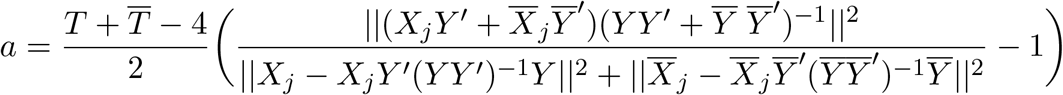

where ||.|| is the Euclidean norm of a vector and I is the regularized incomplete beta function.

Also,

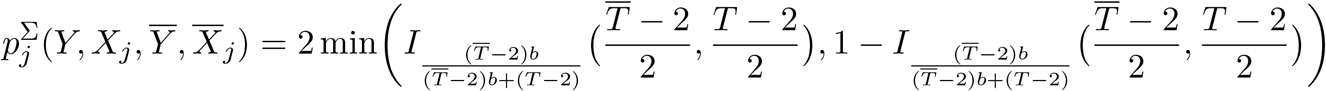

where

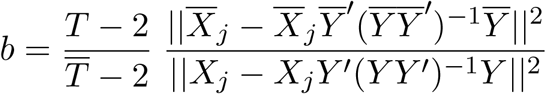

Then the the p-value defined to test 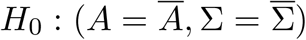is:

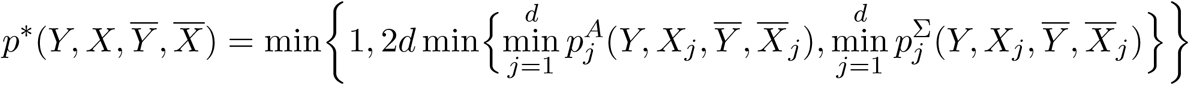

This Bonferroni corrected P-value can be easily shown to be valid for *H*_0_ in the sense that 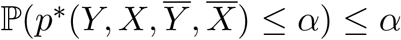 for all α∈ [0, 1] under *H*_0_.

**Remark**: When Σ and 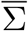 are not diagonal, the Bonferroni correction described above does not have the power to detect whether covariances between error terms among different dimensions are equal or not. In other words, If we have 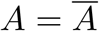and 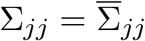for all 1 ≤ *j* ≤ *d*, but the there exist 1 ≤*j ≠ j*^’^ ≤*d* such that 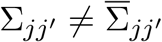, then the offered Bonferroni correction would not reject that.

**Remark**: There are possible scenarios in which the distribution of neural features by itself is not informative enough about the change in the encoding model, unless we have already fixed the movement intention. This is why inspection of model drifts on the conditional distribution ℙ(*X*|*Y*) using the procedure above can work better on those cases.

